# Engineering precursor pools for increasing production of odd-chain fatty acids in *Yarrowia lipolytica*

**DOI:** 10.1101/2020.10.30.362327

**Authors:** Young-Kyoung Park, Florence Bordes, Fabien Letisse, Jean-Marc Nicaud

## Abstract

Microbial production of lipids is one of the promising alternatives to fossil fuels with increasing environmental and energy concern. Odd-chain fatty acids (OCFA), a type of unusual lipids, are recently gaining a lot of interest as target compounds in microbial production due to their diverse applications in the medical, pharmaceutical, and chemical industries. In this study, we aimed to enhance the pool of precursors with three-carbon chain (propionyl-CoA) and five-carbon chain (β-ketovaleryl-CoA) for the production of OCFAs in *Yarrowia lipolytica*. We evaluated different propionate-activating enzymes and the overexpression of propionyl-CoA transferase gene from *Ralstonia eutropha* increased the accumulation of OCFAs by 3.8 times over control strain, indicating propionate activation is the limiting step of OCFAs synthesis. It was shown that acetate supplement was necessary to restore growth and to produce a higher OCFA contents in total lipids, suggesting the balance of the precursors between acetyl-CoA and propionyl-CoA is crucial for OCFA accumulation. To improve β-ketovaleryl-CoA pools for further increase of OCFA production, we co-expressed the *bktB* encoding β-ketothiolase in the producing strain, and the OCFA production was increased by 33 % compared to control. Combining strain engineering and the optimization of the C/N ratio promoted the OCFA production up to 1.87 g/L representing 62% of total lipids, the highest recombinant OCFAs titer reported in yeast, up to date. This study provides a strong basis for the microbial production of OCFAs and its derivatives having high potentials in a wide range of applications.

## 1. Introduction

Microbial production of fuels and green chemicals are considered as a promising alternative to fossil fuels. They offer multiple advantages over plant oils or animal fats such as not competing with food, being less dependent on environmental conditions, and enabling a tunable composition of products. Among several target products from engineered microorganisms, high-value products not readily obtainable through the traditional processes are gaining a lot of interest because they can be produced through an economically feasible process. Odd-chain fatty acids (OCFAs), one of the value-added lipids, are products with potential because they can be used in a variety of applications. For example, *cis*-9-heptadecenoic acid has anti-inflammatory effects and can help treat psoriasis, allergies, and autoimmune diseases [Degwert et al. 1998]. Pentadecanoic acid and heptadecanoic acid can be used as standard compounds of biomarkers for food intake in dietary assessments, the risk of coronary heart disease (CHD), and the risk of type II diabetes mellitus [Jenkins et al. 2015; Forouhi et al. 2014; Pedersen et al. 2016; Pfeuffer and Jaudszus, 2016]. In addition, OCFAs and their derivatives are precursors for manufacturing substances such as pesticides, flavor and fragrance compounds, hydraulic fluids, plasticizers, coatings, and other industrial chemicals [Avis, 2000; Clausen et al. 2010; Köckritz et al. 2010; Fitton and Goa, 1991].

OCFAs are generated by the incorporation of propionyl-CoA in place of acetyl-CoA in the initial condensation step of the fatty acid (FA) synthesis [Ingram et al. 1977; Fulco, 1983]. It is reported that propionyl-CoA can be synthesized from propionate in bacteria and yeast [Ingram et al. 1977; Pronk et al. 1994], and some studies described the production of OCFAs from propionate supplementation without genetic engineering in *Saccharomyces cerevisiae, Yarrowia lipolytica, Cryptococcus curvatus, Rhodococcus* etc. [Kolouchova et al. 2015; Fontanille et al. 2012; Zheng et al. 2012; Bhatia et al. 2019]. Despite a number of applications, studies aiming to produce microbial OCFAs are limited to fermentation with the wild-type strains, resulting in relatively low biomass and oil content [Zhang et al. 2020]. Therefore, the development of platform microorganisms by metabolic engineering for efficient OCFA production is necessary for large-scale application.

Synthesis of adequate precursor levels for the production of target compounds has been considered one of the crucial strategies to improve the productivity of cell factories in metabolic engineering. The importance of the pool sizes was underlined in several studies for the production of desired compounds [Hong et al. 2017]. For example, acetyl-CoA is a precursor of a variety of biotechnology products, including FAs, 1-butanol, polyhydroxyalkanoates, polyketides, and isoprenoids, *etc*. [Nielsen, 2014; Krivoruchko et al. 2015]. A number of studies have focused on increasing acetyl-CoA pools, by engineering the pyruvate dehydrogenase complex (PDH) bypass route for producing amorphadiene in *S. cerevisiae*, or by introducing a cytosolic form of PDH or heterologous PDH for producing 1-butanol, and so on [Shiba et al. 2007; Lian et al. 2014].

The increase of propionyl-CoA pool, a major precursor of OCFAs, was previously reported in the study of producing poly(3-hydroxybutyrate-co-3-hydroxyvalerate) by introducing propionate-activating enzymes (*prpP* and *prpE*) in *Escherichia coli* [Liu et al. 2009]. For the OCFA production, only a few studies demonstrated the improvement of propionyl-CoA pools, so far. Wu and San have shown that the introduction of *prpE* from *Salmonella enterica*, an enzyme catalyzing the ligation of propionate and free CoA to propionyl-CoA, resulted in the improved production of OCFAs in *E. coli* (mainly undecanoic acid (C11:0), tridecanoic acid (C13:0), and pentadecanoic acid (C15:0)) [Wu and San, 2014]. The production of very short odd-chain fuels and chemicals was reported in *E. coli via* overexpression of *pct* gene encoding for a propionyl-CoA transferase from *Megasphaera elsdenii* and further genetic engineering [Tseng and Prather, 2012]. Recently, the improved production of OCFAs by preventing the propionyl-CoA catabolism through the methyl citrate pathway was described in *Y. lipolytica* [Park et al. 2018].

The yeast *Y. lipolytica* is a widely recognized oleaginous yeast presenting high acetyl-CoA flux and an oil sequestration mechanism. Lipid metabolism of this yeast has been thoroughly studied and applied for engineering to improve lipid synthesis mostly even-chain FA (ECFA) or produce valuable FA derivatives such as conjugated linoleic acid, cyclopropane FAs, and cocoa butter-like oils [Imatoukene et al. 2017; Czerwiec et al. 2019; Papanikolaou et al. 2003]. In addition, *Y. lipolytica* can grow in a broad range of substrates *via* native metabolism or synthetic pathway [Ledesma-Amaro and Nicaud, 2016]. The recently developed synthetic biology tools for *Y. lipolytica* also make the yeast more promising chassis for biotechnological production processes [Abdel-Mawgoud et al. 2018; Larroude et al. 2018].

In this study, we aimed to enhance the precursor pools for the production of OCFAs in *Y. lipolytica*. We compared propionate-activating enzymes from different origins to enhance the three-carbon chain precursor pools (C3, propionyl-CoA) and examined OCFA production. Further engineering to improve OCFA production by boosting TAG synthesis and accumulation, and overexpressing *RebktB* which helps to increase five-carbon chain precursor (C5, β-ketovaleryl-CoA) was investigated. Additionally, optimization of the C/N ratio was explored to increase the production of OCFAs.

## 2. Materials and Methods

### 2.1. Strains and media

Media and growth conditions for *E. coli* were previously described by Sambrook and Russell [Sambrook and Russel, 2001]; those for *Y. lipolytica* were previously described by Barth and Gaillardin [Barth and Gaillardin, 1997]. Rich medium (YPD) was prepared containing 1% (w/v) yeast extract, 2% (w/v) peptone, and 2% (w/v) glucose. Minimal glucose medium (YNBD) were prepared containing 0.17% (w/v) yeast nitrogen base (without amino acids and ammonium sulfate, YNBww), 0.5% (w/v) NH_4_Cl, 50 mM KH_2_PO_4_-Na_2_HPO_4_ (pH 6.8), and 2% (w/v) glucose. When minimal media containing different carbon sources and different concentrations were used, the media were named according to the following nomenclature YNBDxPyAz, where D, P and A are glucose, propionate, and acetate, respectively and x,y, and z are the concentration (in %w/v) of these compounds. To complement strain auxotrophy, 0.1 g/L of uracil or leucine was added as necessary. To screen for hygromycin resistance, 250 μg/mL of hygromycin was added to YPD or YNBD media. Solid media were prepared by adding 1.5% (w/v) agar.

### 2.2. Construction of plasmids and strains (*E. coli* and *Y. lipolytica*)

We used standard molecular genetic techniques [Sambrook and Russell, 2001]. Restriction enzymes were obtained from New England Biolabs (Ipswich, MA, USA). PCR amplification was performed in an Eppendorf 2720 Thermal Cycler with either Q5 High-Fidelity DNA Polymerase (New England Biolabs) or GoTaq DNA polymerases (Progema, WI, USA). PCR fragments were purified using a PCR Purification Kit (Macherey-Nagel, Duren, Germany), and plasmids were purified with a Plasmid Miniprep Kit (Macherey-Nagel).

The cassettes of gene expression or disruption were prepared by *NotI* digestion and transformed into *Y. lipolytica* strains using the lithium acetate method, as described previously [Barth and Gaillardin, 1997]. Gene integration and disruption were verified *via* colony PCR using the primers listed in Supplementary Table 1. To construct prototrophic strains, the *LEU2* fragment from plasmid JME2563 or a *URA3* fragment from plasmid JME1046 was transformed. All the strains and plasmids used in this study are listed in Table 1.

**Table 1.**
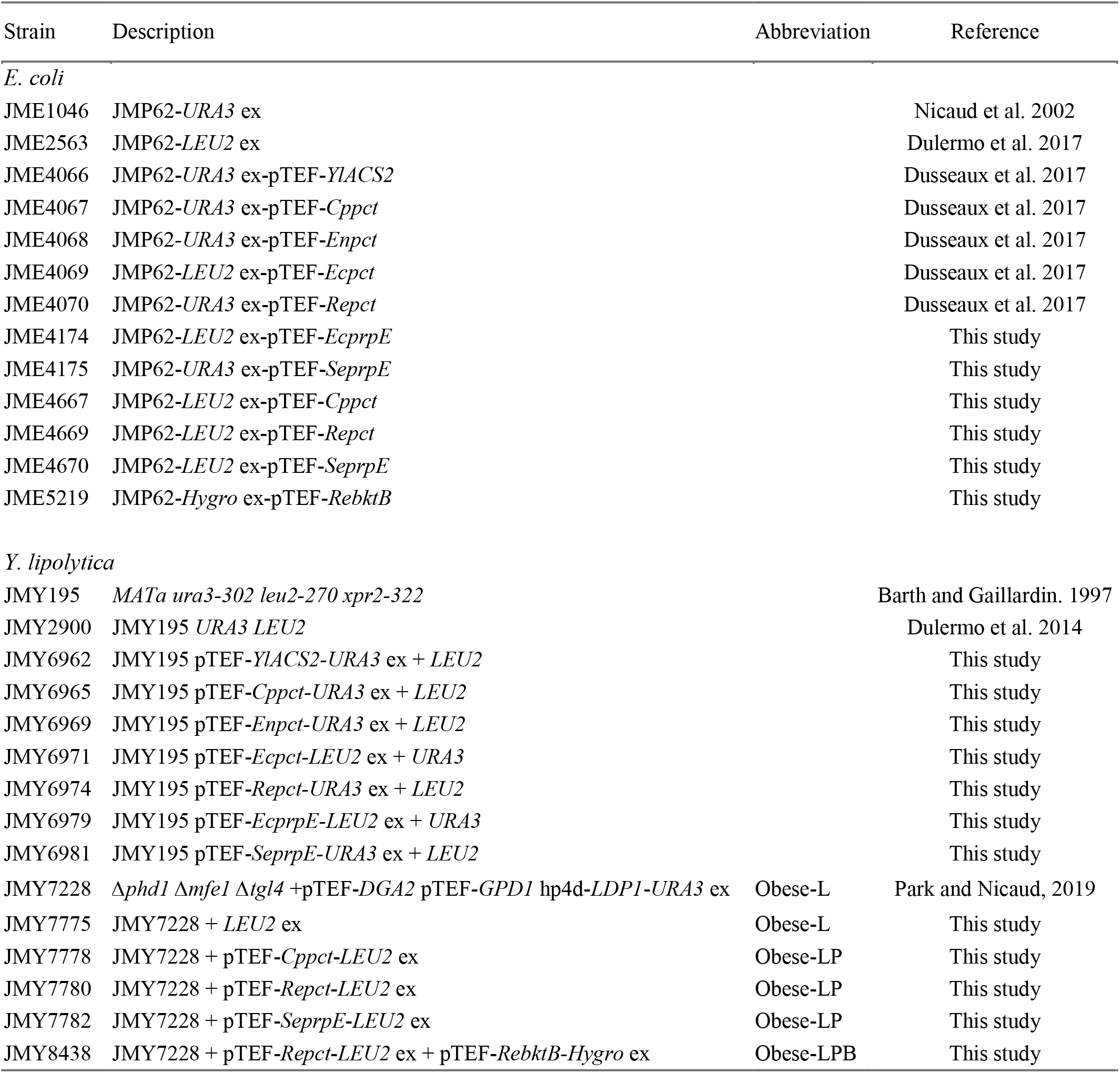
The strains used in this study.

### 2.3. Culture conditions for cell growth

To test growth in liquid culture, pre-cultures were inoculated in the YNBD1 medium and grown overnight (28°C, 180 rpm). The cells were inoculated to fresh YNB medium with an initial OD = 0.1 with different carbon sources depending on the purpose of the experiment. The strains were cultivated at 28°C with constant shaking for 120 hours. Growth was monitored by measuring the OD values every 30 min using a microplate reader (Biotek Synergy MX, Biotek Instruments, Colmar, France). For each strain and set of conditions, we used two biological replicates.

### 2.4. Culture conditions for lipid accumulation

The lipid biosynthesis experiments were carried out in minimal media under nitrogen-limited conditions, and the cultures were prepared as follows: an initial pre-culture was established by inoculating in 10 mL of YPD medium and grown overnight at 28°C and 180 rpm. The cells were washed with sterile distilled water and used inoculating to 50 mL of minimal medium in 250 mL Erlenmeyer flasks containing 0.17% (w/v) yeast nitrogen base (without amino acids and ammonium sulfate, YNBww, Difco), 0.15% (w/v) NH_4_Cl, 50mM KH_2_PO_4_-Na_2_HPO_4_ buffer (pH 6.8). The following carbon sources were added as a sole carbon source or mixture depending on the genotype and the experiment: 0.5 – 4% (w/v) glucose, 0.5 – 2% (w/v) propionate, and 0.5 – 2% (w/v) acetate. The cultures were then incubated at 28 °C and 180 rpm for 5 – 8 days. For each strain and set of conditions, we used two biological replicates.

### 2.5. Substrates determination

The substrates used in this study were identified and quantified by HPLC. Filtered aliquots of the culture medium were analyzed by the UltiMate 3000 system (Thermo Fisher Scientific, UK) using an Aminex HPX-87H column (300 mm × 7.8 mm, Bio-RAD, USA) coupled to UV and RI detectors. Glucose was analyzed by the RI detector, and propionate and acetate were analyzed by UV detectors (210 nm). The mobile phase used was 0.01 N H2SO4 with a flow rate of 0.6 mL/min and the column temperature was T = 35 °C. Identification and quantification were achieved *via* comparisons to standard compounds. For each data point, we used at least two biological replicates and calculated average and standard deviation values.

### 2.6. Lipid determination

Lipids were extracted from 10 – 20 mg of freeze-dried cells and converted into FA methyl esters (FAMEs) using the procedure described by Browse et al. [Browse et al. 1986]. The FAMEs were then analyzed using gas chromatography (GC), which was carried out with a Varian 3900 instrument (Varian Inc. USA) equipped with a flame ionization detector and a Varian FactorFour vf-23ms column, where the bleed specification at 260 °C is 3 pA (30 m, 0.25 mm, 0.25 μm). The FAMEs were identified *via* comparisons with commercial standards (FAME32, Supelco) and quantified using the internal standard, 100 μg of commercial dodecanoic acid (Sigma-Aldrich, USA). Commercial standards of OCFAs (Odd Carbon Straight Chains Kit containing 9 FAs, OC9, Merck, Germany) were converted into their FAMEs using the same method employed with the yeast samples. They were then analyzed by GC to identify the OCFAs from the yeast samples. For each data point, we used at least two biological replicates and calculated average and standard deviation values.

To determine the dry cell weight (DCW), 2 mL of the culture was taken from the flasks, washed, and lysophilized in a pre-weighed tube. The differences in mass corresponded to the mg of cells found in 2 mL of culture.

## 3. Results

### 3.1. Evaluating propionate-activating enzymes for OCFA production

Previous studies described the production of OCFAs with propionate supplementation in wild-type *Y. lipolytica*, which indicated the presence of an endogenous propionyl-CoA synthetase activity in *Y. lipolytica* [Fontanille et al. 2012; Park et al. 2018]. We evaluated heterologous and native enzymes (CoA transferase and CoA synthetase) involved in propionate activation to see if they could improve propionyl-CoA availability for the production of OCFAs (Figure 1 (a)).

**Figure 1.**
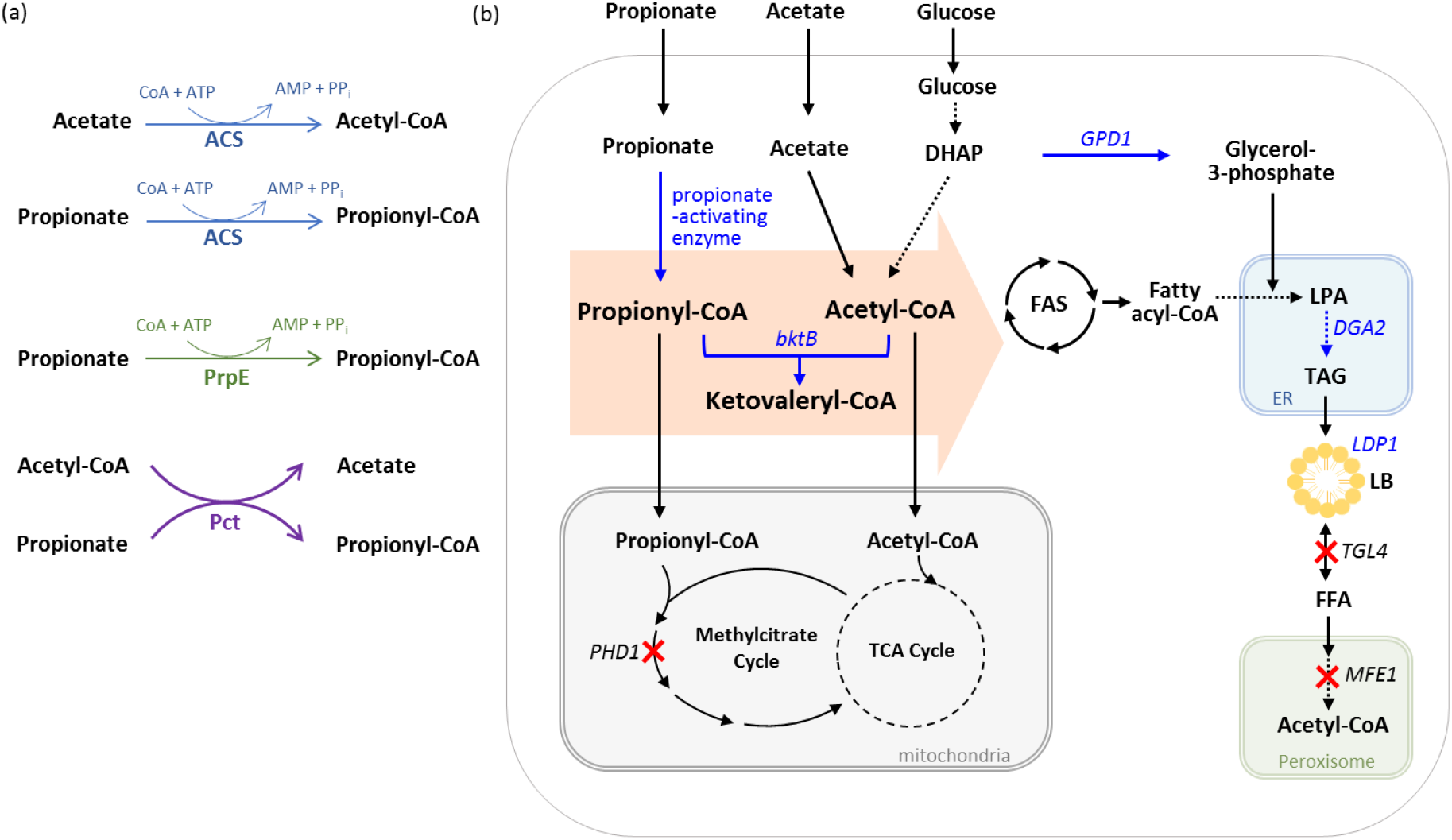
Overview of synthesis of lipids including OCFAs in *Y. lipolytica*. (a) Reaction of propionate activating enzymes used in this study, (b) overall strain engineering for OCFA production applied in this study. Engineered steps by overexpression are indicated by blue arrows and the corresponding genes are written in blue. Inactivated steps are indicated by red cross and the corresponding genes are written in black. The precursors of OCFAs are grouped in orange arrow. Multiple steps are shown as dashed arrows. *bktB*, β-ketothiolase; *GPD1*, glycerol-3-phosphate dehydrogenase; *DGA2*, acyl-CoA:diacylglycerol acyltransferase; *LDP1*, lipid droplet protein; *PHD1*, 2-methylcitrate dehydratase; *TGL4*, triglyceride lipase; *MFE1*, multifunctional enzyme; DHAP, dihydroxyacetone phosphate; LB, lipid body.

Pct (propionyl-CoA transferase) catalyzes the transfer reaction of the CoA moiety from the donor (generally acetyl-CoA) to the acceptor through ping-pong mechanism; the range of substrate is broad and diverse depending on the origin [Volodina et al. 2014]. PrpE (propionyl-CoA synthetase) and Acs (acetyl-CoA synthetase) are both acyl-CoA synthetase enzymes (EC 6.2.1) and share sequence similarities as well as common reaction features. PrpE (propionyl-CoA synthetase) is a bacterial propionate-activating enzyme, the overexpression of *prpE* from *Salmonella enterica* showed the increase of propionate activation in *E. coli* [Wu and San, 2014]. In yeast, it is known that acetyl-CoA synthetase can catalyze the formation of propionyl-CoA from propionate and coenzyme A [Jones and Lipmann, 1955; Pronk et al. 1994], *Y. lipolytica* has one gene (*ACS2*, *YALI0F05962g)* encoding acetyl-CoA synthetase while there are two genes (*ACS1, ACS2*) in *S. cerevisiae*. Seven genes, *pct* (from *Ralstonia eutropha, Clostridium propionicum, E. coli*, and *Emiricella nidulans), prpE* (from *E. coli* and *S. enterica)* and native *ACS2*, were evaluated in *Y. lipolytica* wild-type strain. All genes were synthesized with codon optimization for *Y. lipolytica* and expressed under the constitutive pTEF promoter. The sequences of the gene used in this study are described in Supplementary Table 2.

*Y. lipolytica* strains expressing individual propionate activating enzymes were grown on minimal medium with various carbon sources (Figure 2, Supplementary Figure 1 and 2). The growth of all strains on glucose in YNBD0.5 was mostly similar, while the growth on propionate in YNBP0.5 highly depended on the gene being overexpressed (Supplementary Figure 1). On propionate as a sole carbon source *Enpct*-, *Ecpct*-, *EcprpE*- and *SeprpE-* expressing strains grew similarly to wild-type (Supplementary Figure 1 (b)), whereas three strains showed impaired growth, the *Cppct-, Repct-* and *ACS2*-expressing strains (Figure 2 and Table 2). The overexpression of native *ACS2* resulted in a decrease of growth by 30% and 25% in final OD and growth rate, respectively. In case of *Cppct*-expressing strain, the growth was slower than wild-type, but reached similar final OD. The growth of *Repct*-expressing strain was the most affected on propionate as it grew much slower than *Cppct*-expressing strain (Figure 2 and Supplementary figure 2). The growth rate was decreased by 54% compared to wild-type (μ_max_ *Repct* = 0.057 h^-1^ *vs*. μ_max_ wild-type = 0.123 h^-1^), and the final OD was lower than wild-type by 22%. The lower growth of *Cppct-* and *Repct*-expressing strains compared to wild-type could have different explanations, as the reaction consumes acetyl-CoA and propionate and produces acetate and propionyl-CoA, it could be due to the low acetyl-CoA pool for the growth or the toxicity of released acetate or propionyl-CoA.

**Figure 2.**
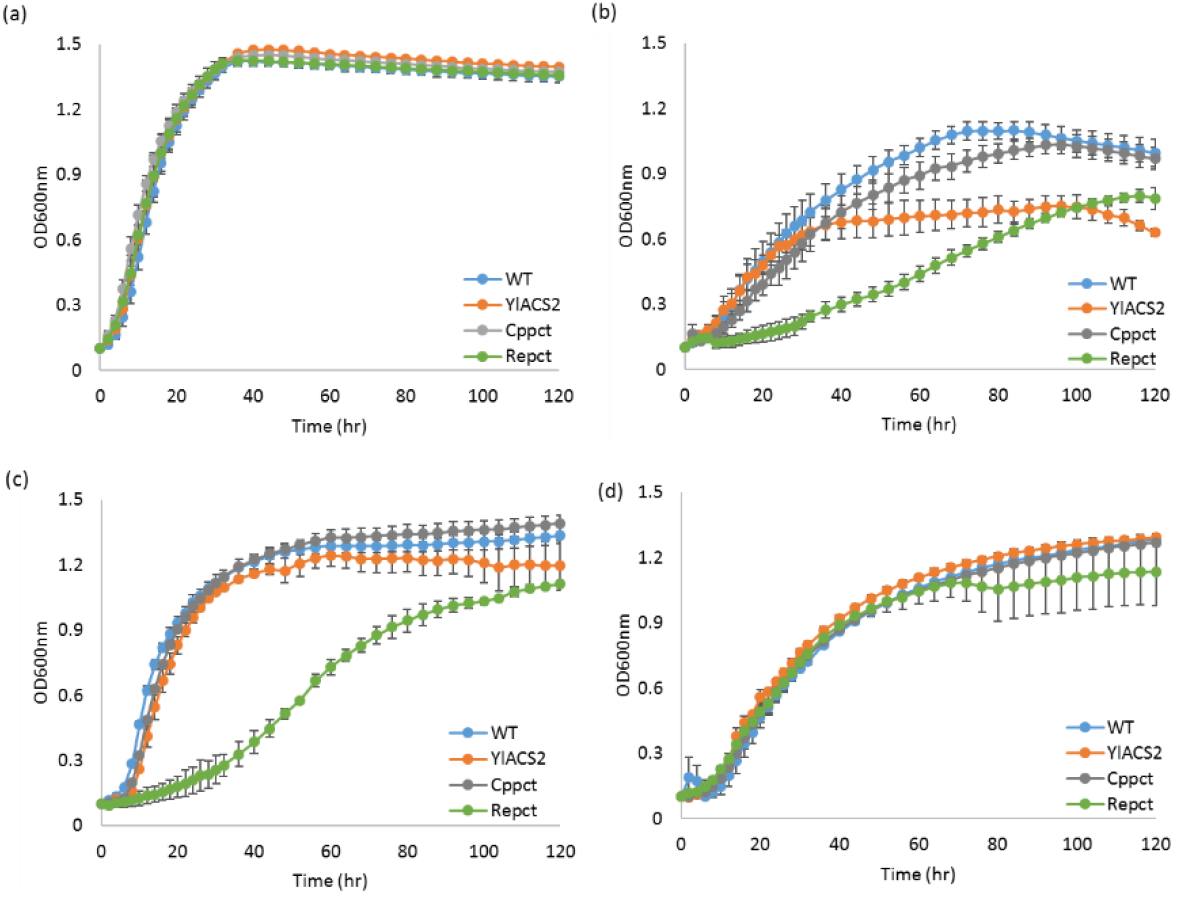
Comparison of growth in wild-type and strains overexpressing *YlACS2*, *Cppct*, and *Repct* genes with different substrates. (a) YNBD0.5, (b) YNBP0.5, (c) YNBD0.5P1, and (d) YNBP1A0.5. Averages were obtained from two replicate experiments.

**Table 2.** Maximum growth rate (μ_max_, h^-1^) of the strains overexpressing propionate activating genes. Strains were grown in YNBD0.5, YNBP0.5, YNBD0.5P1, and YNBP1A0.5, respectively. Average values were obtained from two replicate experiments.

|  | YNBD0.5 | YNBP0.5 | YNBD0.5P1 | YNBP1A0.5 |
| --- | --- | --- | --- | --- |
| WT | $0.205 \pm 0.010$ | $0.123 \pm 0.043$ | $0.271 \pm 0.018$ | $0.168 \pm 0.011$ |
| *YlACS2* | $0.242 \pm 0.013$ | $0.093 \pm 0.006$ | $0.279 \pm 0.032$ | $0.176 \pm 0.017$ |
| *Cppct* | $0.272 \pm 0.014$ | $0.098 \pm 0.011$ | $0.260 \pm 0.002$ | $0.130 \pm 0.001$ |
| *Enpct* | $0.206 \pm 0.003$ | $0.135 \pm 0.012$ | $0.246 \pm 0.005$ | $0.186 \pm 0.005$ |
| *Ecpct* | $0.249 \pm 0.021$ | $0.126 \pm 0.004$ | $0.278 \pm 0.001$ | $0.144 \pm 0.008$ |
| *Repct* | $0.262 \pm 0.006$ | $0.057 \pm 0.015$ | $0.055 \pm 0.002$ | $0.141 \pm 0.023$ |
| *EcprpE* | $0.238 \pm 0.009$ | $0.093 \pm 0.011$ | $0.279 \pm 0.014$ | $0.141 \pm 0.011$ |
| *SeprpE* | $0.181 \pm 0.010$ | $0.095 \pm 0.003$ | $0.181 \pm 0.006$ | $0.105 \pm 0.004$ |

In order to verify this, we tested cell growth on propionate with glucose or acetate as co-substrate (Figure 2 and Supplementary Figure 1). When glucose was added to the propionate medium, the growth of the *Cppct*-expressing strain was restored. For *ACS2-*expressing strain, the final OD was almost completely recovered compared to wild-type (89.5% of the final OD). The *Repct*-expressing strain remained the most impacted, even though the addition of glucose slightly improved growth compared to the one on propionate as a sole carbon source, the growth rate is still 80% lower than the one of the wild-type (μ_max_ *Repct* = 0.055 h^-1^ *vs*. μ_max_ wild-type = 0.271 h^-1^), and still showed a slightly lower final OD than wild-type. Thus, the addition of glucose to the propionate medium helped the cell growth but differentially depending on the strain. When acetate is added to propionate instead of glucose, all strains showed similar growth, including *Repct*-expressing strain (Figure 2 (d)).

In order to see if the overexpression of propionate-activating genes improves the accumulation of OCFAs, the strains were cultivated in nitrogen-limited conditions (0.15% (w/v) NH_4_Cl) with different substrates and the lipid accumulation was quantified. The OCFA accumulation of the strains was also variable depending on the substrate as it was observed for cell growth (Figure 3). The addition of glucose and/or acetate in propionate medium resulted in a decrease of OCFA accumulation in wild-type strain. The overexpression of native *ACS2* exhibited a negative effect on OCFAs accumulation compared to wild-type in all conditions. The overexpression of *Cppct, Repct*, and *SeprpE* showed an increase of OCFAs (% of total FAs) compared to control for all culture conditions only except for *Repct*-expressing strain in YNBD0.5P1. When we added acetate to YNBD0.5P1, the ratio of OCFAs to total FAs decreased in all strains including wild-type except for *Cppct*-, *Repct*-, and *SeprpE-*expressing strain. Especially, the *Repct*-expressing strain accumulated the highest ratio of OCFAs to total FAs, 53.2%, in this experiment. Therefore, the three genes (*Cppct*, *Repct*, and *SeprpE*) showing a significant improvement of OCFAs were selected for subsequent strain engineering.

**Figure 3.**
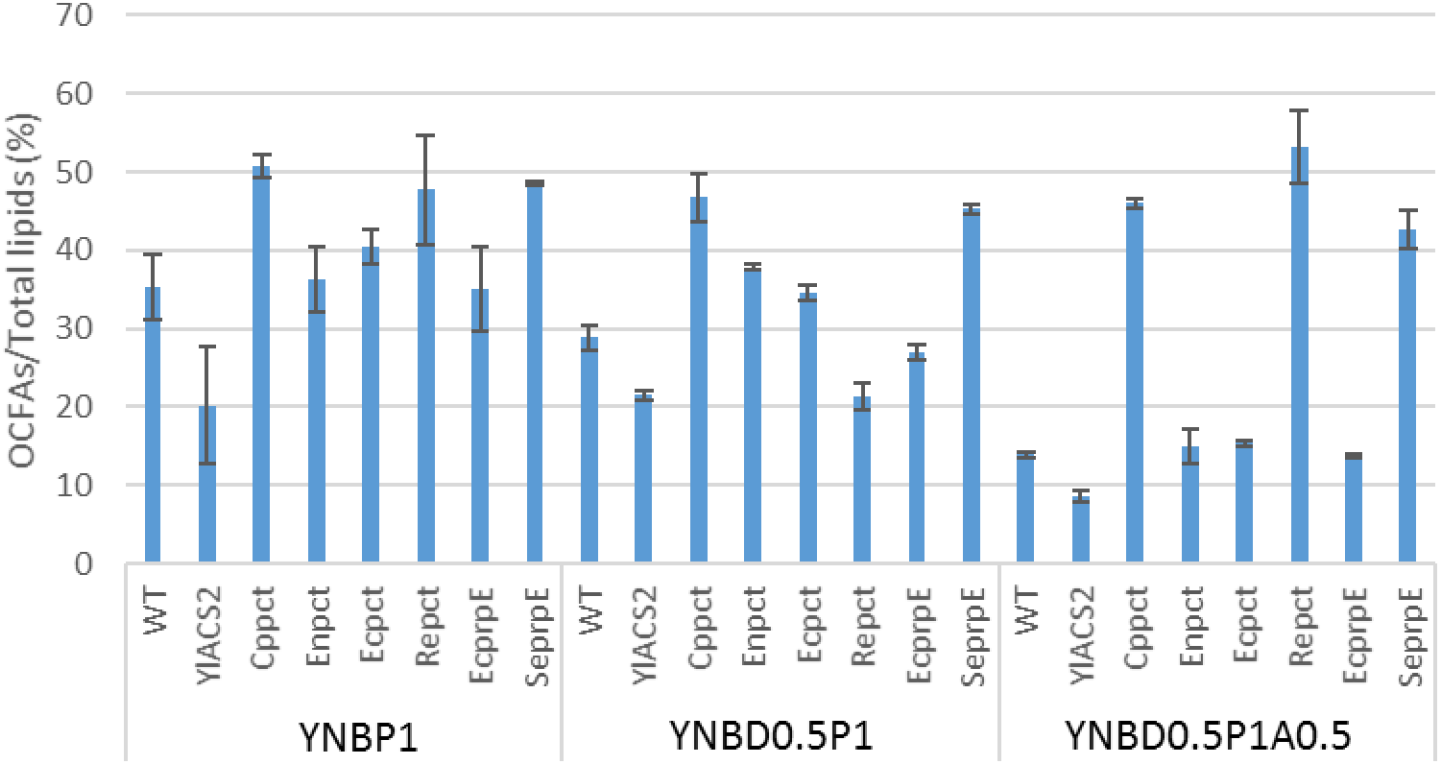
Accumulation of OCFAs in *Y. lipolytica* strains overexpressing propionate activating genes (*ACS*, *pct*, and *prpE)*. The strains were cultivated in YNBP1, YNBD0.5P1, and YNBD0.5P1A0.5 medium for 120 hours. Averages and standard deviations were obtained from two replicate experiments.

### 3.2. Improving OCFA production by engineering obese strains

We previously engineered *Y. lipolytica* to accumulate a high amount of OCFAs, 41.9% of total lipids [Park et al. 2018; Park and Nicaud, 2019]. One of the main modifications was the deletion of the *PHD1* gene preventing propionyl-CoA consumption through the methyl citrate pathway. Through further modifications, deletion of *MFE1* to inhibit β-oxidation and *TGL4* to inhibit triacylglycerols (TAG) remobilization, and overexpression of *GPD1* and *DGA2* to push and pull TAG biosynthesis, we constructed obese strain (JMY3776). In addition, we overexpressed the lipid droplet protein *LDP1* to enhance the storage of TAG and named the strain as obese-L strain (JMY7228) [Bhutada et al. 2018; Park and Nicaud, 2019]. All these modifications are described in Figure 1(b) together with newly introduced modifications in this study.

The three selected genes (*Cppct, Repct*, and *SeprpE*) were separately overexpressed in obese-L strain, obtaining obese-LP strains. To assess the lipid accumulation of the strains, flask culture was performed in YNBD2P0.5A1 (2% (w/v) glucose, 0.5% (w/v) propionate, 1% (w/v) acetate with a C/N ratio = 30) (Figure 4). All obese-LP strains accumulated OCFAs more than 54% of total FAs (Figure 4 (b)). The obese-LP (*Repct*) strain exhibited the highest amount of OCFAs, 61.7% of total lipids. Whereas the obese-L strain (Figure 4 (a) and (c)) accumulates a majority of ECFAs (16:0 and C18:1; palmitic acid and oleic acid, respectively), C17:1 (heptadecenoic acid) was the most abundant OCFA in all the obese-LP strains consistently with our previous study (Figure 4 (d)). The highest ratio of C17:1 to total lipids (45.6%) was obtained in obese-LP (*SeprpE*) strain. The second major FA from obese-LP strain was C18:1 (between 15% and 20% depending on the strain), and other FAs are lower than 10 % of total FAs (Supplementary Figure 3).

**Figure 4.**
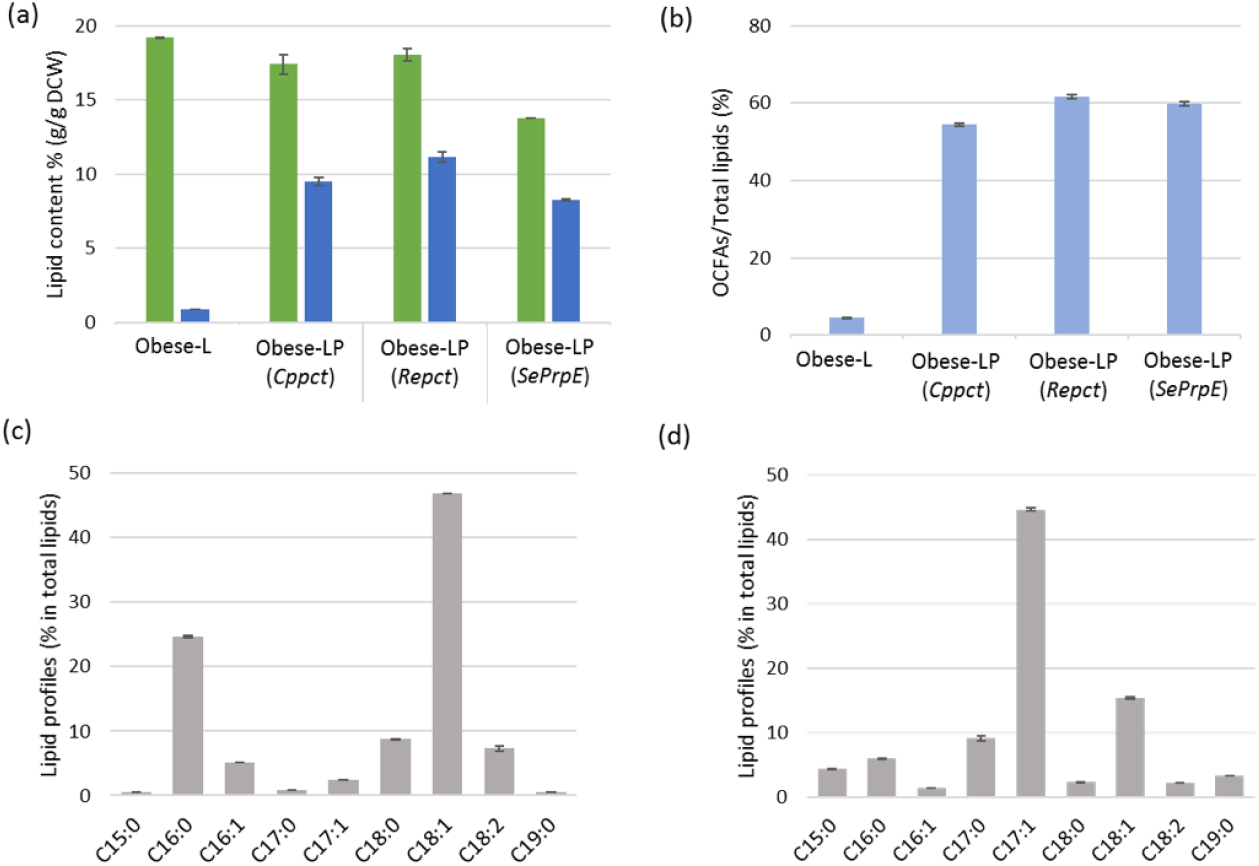
Lipid accumulation of obese strains expressing *pct* and *prpE* genes. (a) The percentage of FAs in the DCW, total FAs are indicated in green and OCFAs are in blue, (b) the ratio of OCFAs to total lipids (%), (c) lipid profiles (% of total lipids) of obese-L strain, (d) the lipid profiles (% of total lipids) of obese-LP (*Repct*) strain. The strains were cultivated in the YNBD2P0.5A1 medium for 120 hours. Averages and standard deviations were obtained from two replicate experiments.

All three obese-LP strains showed a higher propionate consumption compared to the control strain at the beginning of the culture (until 50 hours) (Supplementary Figure 4). Among them, the obese-LP (*SeprpE*) strain consumed the highest amount of propionate, 2.4 g/L, at a constant rate. The obese-LP (*Repct*) and the obese-LP (*Cppct*) strains also utilized more propionate (1.8 g/L, 1.2 g/L, respectively) than the control strain (1.1 g/L).

### 3.3. Supplementing acetate in OCFA production

We assume that the addition of acetate increases acetyl-CoA pools and improves the level of ECFA at the expense of OCFA. In order to investigate how the addition of acetate has an impact on the ratio of OCFA and ECFA, we first cultured the obese-L strain with and without acetate (YNBD1P0.5 and YNBD1P0.5A1, respectively). The ratio of OCFA to total lipids was significantly dropped from 28.9% to 3.8% when acetate (1%) was added to the medium (Supplementary Table 3). We then explored the lipid accumulation with different amounts of acetate to identify the optimal ratio of propionate and acetate for OCFA production. The obese-LP (*Repct*) strain was used for this test, glucose was also added to provide a carbon source for biomass as the previous study showed a beneficial effect of this strategy [Park et al. 2018]. Different ratios in the concentration of propionate and acetate (from 2:1 to 1:4) were applied with a constant propionate concentration (0.5% (w/v)), and glucose was adjusted to meet the same total carbon amount in all conditions. The more acetate was added, the lower ratio of OCFAs to total FAs (from 66.3 to 50.7 %) was obtained as expected (Figure 5). Heptadecenoic acid (C17:1) and nonadecanoic acid (C19:0) were decreased, while oleic acid (C18:1) was increased with increasing acetate amount. The other OCFAs, pentadecanoic acid (C15:0) and heptadecanoic acid (C17:0), showed relatively steady levels regardless of acetate amount. The highest concentration of acetate resulted in the highest biomass and total lipid content (%, g/g DCW). The combination of 0.5% (w/v) propionate and 1% (w/v) acetate was selected as the best combination for OCFA production regarding the total lipid content and the ratio of OCFA in total lipids.

**Figure 5.**
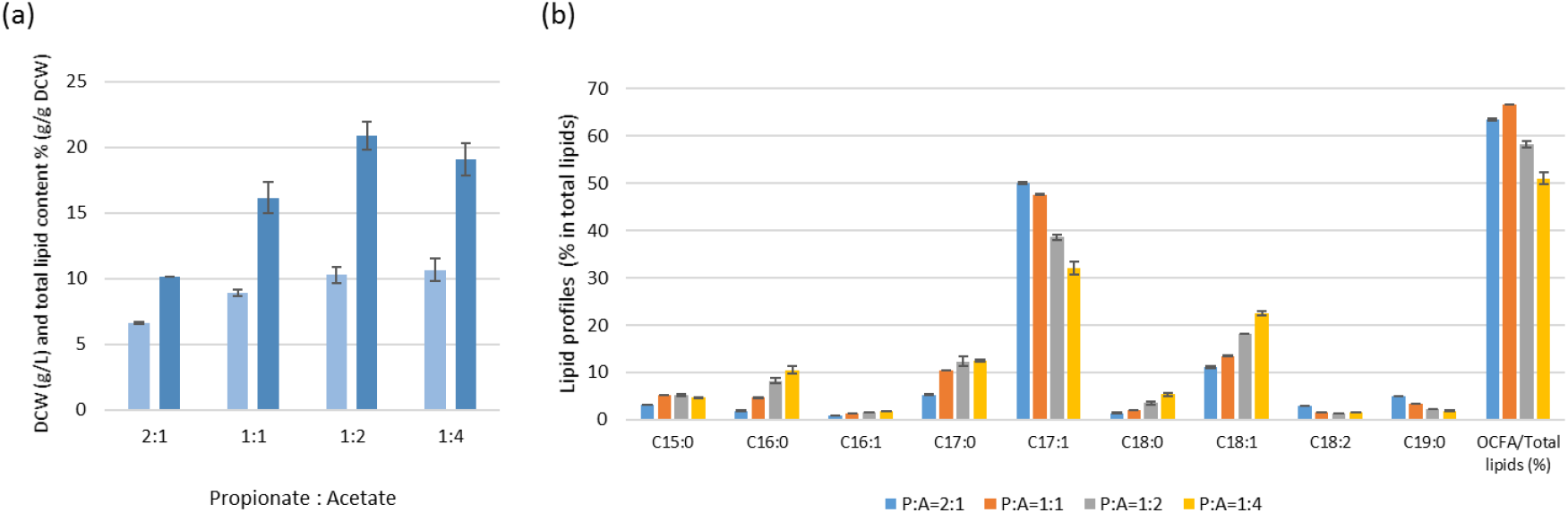
Lipid accumulation of JMY7780 (obese-LP (*Repct)*) grown with the different ratio of propionate and acetate. (a) DCW (g/L) in light blue and total lipid content (%, g/g DCW) in blue, (b) the lipid profiles (% of total lipids). The strains were cultivated for 120 hours. Averages and standard deviations were obtained from two replicate experiments.

### 3.4. Increasing C5 precursor pools by overexpressing *bktB*

To further improve precursor pools of OCFAs, lipid accumulation from five-carbon chain compounds was explored. It is reported that valerate (pentanoate, C5) can be utilized as a carbon source for biomass and OCFA production in *Y. lipolytica* [Chakraborty, 2015]. It is explained that the activated form of valerate (valeryl-CoA) enters the β-oxidation to produce acetyl-CoA and propionyl-CoA, resulting in the accumulation of OCFAs. However, the β-oxidation pathway of the obese-LP strain is blocked by *MFE1* deletion, thus the cleavage of C5 compounds to C2 and C3 compounds is inhibited in our strain. We investigated whether *de novo* synthesis of the C5 precursor, β-ketovaleryl-CoA could improve the availability of precursor pools for OCFA production instead of valerate supplementation.

β-Ketothiolase catalyzes the condensation of acetyl-CoA and propionyl-CoA to form β-ketovaleryl-CoA. This enzyme encoded by the *bktB* gene has been used for the synthesis of polyhydroxybutyrate and poly(3-hydroxybutyrate-co-3-hydroxyvalerate) (PHBV) [Mitsky et al. 1998; Tseng and Prather, 2012]. The *bktB* gene from *R. eutropha*, well known as a representative microorganism of the PHBV producer, was selected as a candidate enzyme for synthesizing β-ketovaleryl-CoA in *Y. lipolytica*. The sequence of *bktB* from *R. eutropha* H16 was codon-optimized to *Y. lipolytica* and overexpressed under the constitutive pTEF promoter. To evaluate the effect of *bktB* overexpression on OCFA production, the obese-LP (*Repct*) and obese-LPB (*Repct-RebktB*) strains were grown in YNBD2P0.5A1 medium for 7 days. The production of OCFAs was increased by 33% from 0.99 g/L to 1.32 g/L by overexpressing *bktB* (Figure 6). Total lipid production was also increased by 36% which resulted in a similar ratio of OCFAs to total lipids between obese-LP and obese-LPB strains.

**Figure 6.**
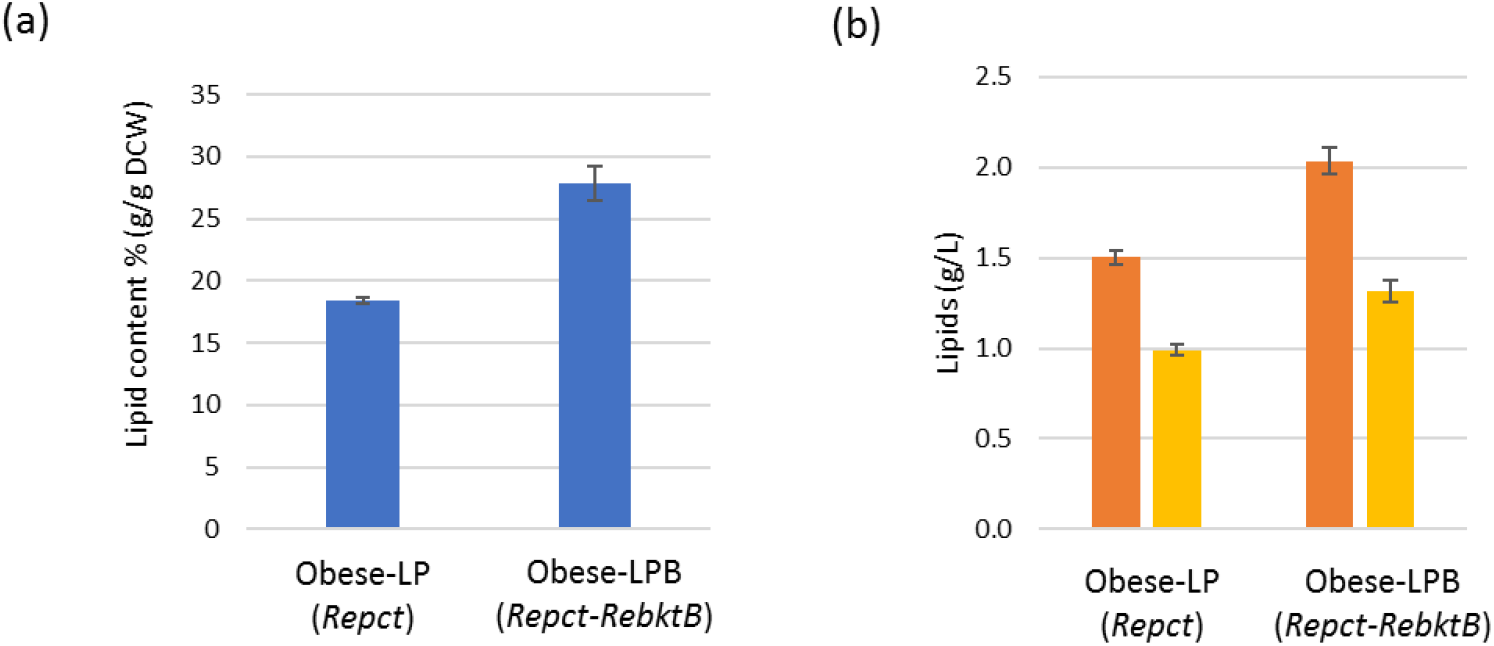
Lipid accumulation of obese-LP (*Repct*) strains with/without overexpression of *RebktB*. (a) The percentage of FAs in the DCW, (b) lipid titer (g/L), total lipids are indicated in orange and OCFAs are in yellow. The strains were cultivated for 168 hours. Averages and standard deviations were obtained from two replicate experiments.

### 3.5. Optimizing C/N ratio

Because the C/N ratio significantly affects triggering lipid accumulation in *Y. lipolytica*, nitrogen-limited media with a high C/N ratio have been used to achieve high levels of lipid accumulation in *Y. lipolytica* [Beopoulos et al. 2009; Lazar et al. 2014; Blazeck et al. 2014]. It is known that the optimal C/N ratio is variable depending on the carbon source and the genotype of the strain [Lazar et al. 2015; Ledesma-Amaro et al. 2016]. The effect of C/N ratios on OCFA productions in the obese-LPB strain was investigated (Figure 7). Different initial C/N ratios were achieved by adjusting NH_4_Cl concentration while holding the initial carbon source concentration constantly (YNBD4P1A2). The ratio of OCFAs to total lipids seems independent of the value of the C/N ratio being maintained at around 62% for all conditions tested. The best lipid accumulation were obtained at the ratio of 45 or 60 depending on the culture time and the optimal C/N ratio of our strain for OCFA production was 45 at 192 hours of culture which produced 1.87 g/L of OCFAs (Figure 7 (b)), the highest OCFA titer obtained from yeasts, so far (Supplementary Table 4).

**Figure 7.**
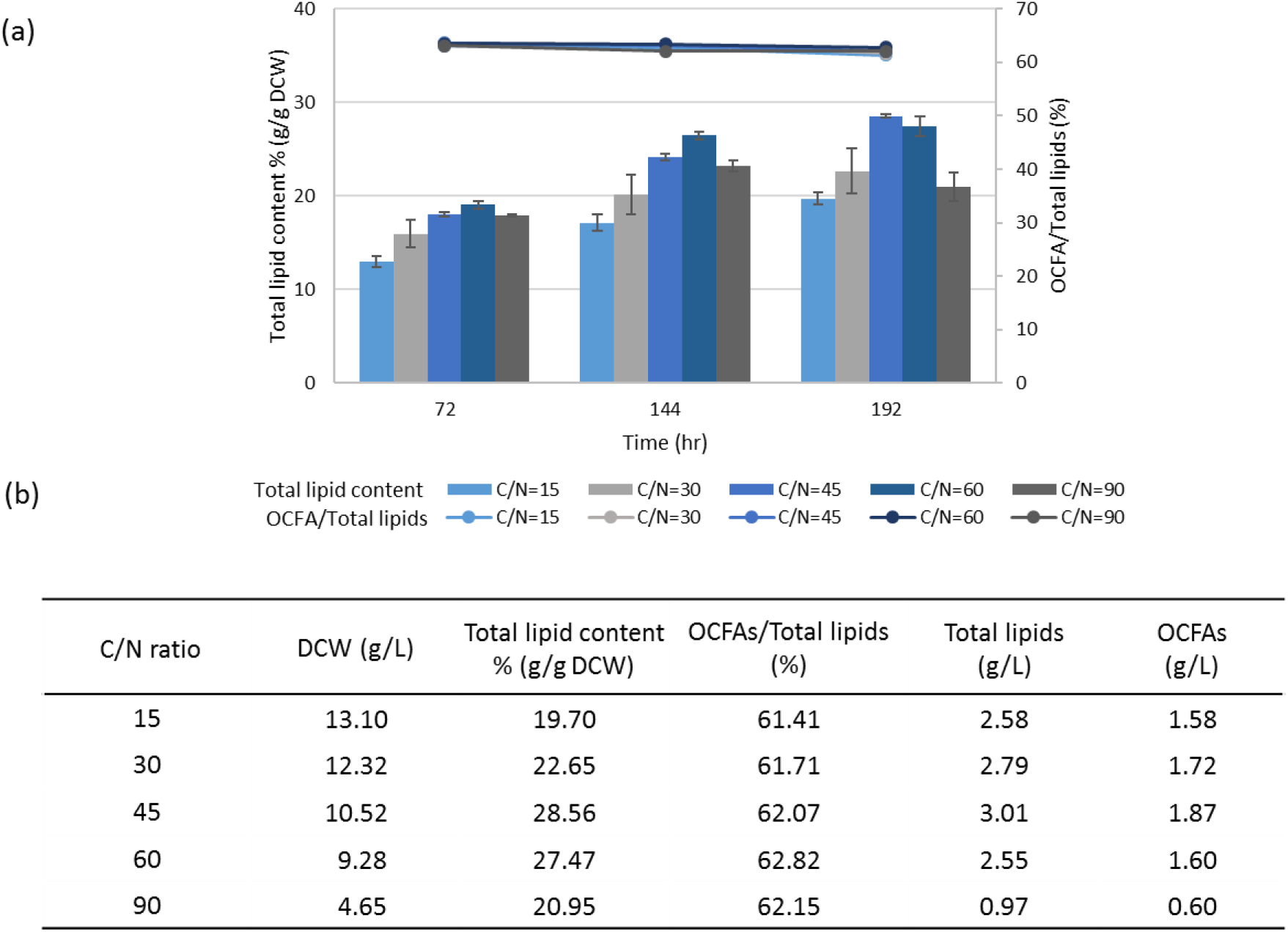
Lipid production by JMY8438 (obese-LPB (*Repct*-*RebktB*)) strain with different C/N ratios. (a) The total lipid content (%, g/g DCW) represented by bar and the ratio of OCFAs in total lipids (%) represented by line, (b) table of lipids and DCW (g/L) at 192 hours of cultivation. Different C/N ratios were obtained by adjusting NH_4_Cl concentration. Averages and standard deviations were obtained from two replicate experiments.

## 4. Discussion

The production of OCFA versus classical ECFA depends only on the first cycle of FA elongation, in which cytosolic propionyl-CoA or acetyl-CoA are respectively condensed with malonyl-CoA. For the OCFA initiation step, it is thus necessary to have a relatively large propionyl-CoA pool compared to acetyl-CoA. But for the subsequent elongation step, the high availability of acetyl-CoA is necessary as the substrate of malonyl-CoA building block. In this study, we thus faced with the dual role of acetyl-CoA for the synthesis of OCFA and discussed our result in the light of this consideration.

As the synthesis of OCFAs is started from propionyl-CoA, can be synthesized from propionate, we first investigated how propionate activating enzymes of different origins could improve the propionyl-CoA availability in cytosol, and therefore, influence the synthesis of the OCFA. Acetyl-CoA synthetases of different origins have shown a promiscuous propionyl-CoA synthetase activity [Luong et al. 2000; Ingram-Smith and Smith, 2006]. However, the detailed studies on the activity of Acs enzyme on propionate or identification of other propionate-activating enzymes in yeast are still limited. In this study, native *Y. lipolytica ACS2* was overexpressed to examine its effect on OCFA production. Its overexpression exhibited a strong negative effect on OCFA accumulation for any substrate compositions in our study, indicating the reaction catalyzed by Acs has a stronger preference for acetate than propionate similar to Acs1p in *S. cerevisiae* [Van den Gerg et al. 1996], thus resulting in a higher amount of ECFAs than OCFAs. The result suggested the further engineering of Acs such as altering substrate specificity by mutagenesis of carboxylate binding pocket [Sofeo et al. 2019] or exchanging native Acs2p with heterologous propionyl-CoA synthetase [Hitschler et al. 2020] might be helpful to increase propionyl-CoA/acetyl-CoA ratio.

Two *prpE* genes (propionyl-CoA synthetase) from *E. coli* and *S. enterica* were overexpressed. The overexpression of *EcprpE* did not show any difference neither in growth nor in OCFA production compared to wild-type, suggesting an inactive or low active enzyme. A higher OCFA production from *SeprpE*-overexpressing strain than wild-type was obtained in all substrate conditions used in this study, showing that our strategy of increasing the propionyl-CoA pool for increasing OCFA production was relevant. Heterologous expression of *prpE* has been a strategy in several studies on propionyl −CoA mediated production, but the effect on target production is various depending on the origin of *prpE* and chassis microorganisms [Wu and San, 2014; Tseng and Prather, 2012; Liu et al. 2009; Callari et al. 2018; Hitschler et al. 2020]. Therefore, the verification and optimization of heterologous *prpE* expression is necessary in each target and platform strain.

Another class of propionate activating enzyme was tested in this study, Pct (propionyl-CoA transferase) catalyzing the transfer of CoA moiety from the acetyl-CoA (donor) to the propionate (acceptor). The *pct*-expressing strains showed different growth on propionate. On this medium, the proportion of OCFA in total lipids of *Ecpct-* and *Enpct-*expressing strains is comparable with the one of the wild-type, whereas the *Cppct-* and *Repct-*expressing strains presented a better accumulation of OCFA than wild-type despite slower growth. We assume that the negative effect on growth for *Cppct*-and *Repct-*expressing strains might be due to the higher Pct activity that led to lower cytosolic acetyl-CoA pool. This correlates with the observed higher accumulation of OCFA as a result of improved propionyl-CoA pool. In order to verify our hypothesis, supplementation of glucose or acetate as an acetyl-CoA supplier on the propionate-containing medium was assessed. The total recovery of growth in the *Repct*-expressing strain was possible only with acetate supplementation, while the growth of the *Cppct*-expressing strain was recovered similar to wild-type in both media. It suggests that the supply of acetyl-CoA from glucose is still limited in *Repct*-expressing strain, representing the highest Pct activity among Pct enzymes tested in this study as confirmed by the highest OCFA accumulation (53.2% of OCFAs in total FAs) on propionate medium supplemented with acetate. The effect of acetate supplementation on OCFA production was then explored in different strains. In non-obese strain including wild-type, the ratio of OCFAs to total FAs was decreased when we added 1% acetate in all strains except for *Repct*-expressing strains. This effect was even more pronounced in the Obese-L strain in which the addition of 1% acetate resulted in only 3.8% of OCFA of total lipids. For the *Repct*-expressing strain, the best compromised condition for cells growth, lipids content, and lipid composition was 2% (w/v) glucose, 0.5% (w/v) propionate and 1% (w/v) acetate. These results indicate the importance of balancing two precursor pools, propionyl-CoA and acetyl-CoA for OCFA production. It also has been reported that the propionyl-CoA/acetyl-CoA ratio affected on the production of angelyl-CoA and 3-ethylphenol which need two precursors together, though it seems difficult to determine a suitable balance of propionyl-CoA and acetyl-CoA concentrations [Callari et al. 2018; Hitschler et al. 2020]. Thus, balancing the precursor pools through optimizing substrates ratio could be one of options to improve these relevant studies.

Using acetate as a substrate has been described in several studies taking advantage of its relatively shorter conversion pathway to acetyl-CoA than general substrates like glucose. The lipid production by utilizing acetate has been reported in oleaginous yeasts such as *Y. lipolytica, Rhodosporidium toruloides*, and *Lipomyces starkeyi* [Fontanille et al. 2012; Xu et al. 2017; Huang et al. 2016; Xavier et al. 2017]. Acetate as well as propionate, substrates used in this study, are major components of volatile fatty acids (VFAs) being easily obtainable from agro-industrial wastes or biodegradable organic wastes. VFAs as a substrate for microbial oil production has recently been gaining a lot of interest due to its low cost [Liu et al. 2017; Llamas et al. 2019]. Thus, the utilization of VFAs for the production of OCFAs in *Y. lipolytica* will be a feasible and sustainable strategy for the scale-up process. To maximize biomass and OCFA production from *Y. lipolytica*, the fermentation conditions like medium compositions, feeding substrates including VFAs, C/N ratio, and pH need to be optimized.

As an alternative to acetate supplementation, the pool of acetyl-CoA can be enhanced by metabolic engineering approach. Because of the importance of acetyl-CoA as a crucial precursor of various products such as isoprenoids, alcohols, sterols, polyketides, polyphenols, and so on, several different strategies have been described, mostly in *S. cerevisiae* [Nielsen, 2014]. Therefore, increasing acetyl-CoA pools by metabolic engineering will be explored in *Y. lipolytica* in a future study which will allow simpler production processes.

An original strategy consisting in the supply of the C5 precursor, β-ketovaleryl-CoA, was investigated, to determine if it can be incorporated into lipid synthesis. β-Ketothiolase encoded by *bktB* from *R. euthropha*, which has been used for the synthesis of PHBV, was selected because of its activity on propionyl-CoA [Mitsky et al. 1998; Tseng and Prather, 2002; Yang et al. 2012]. This strategy has never been explored to improve the OCFA production and the co-expression of *Repct* and *RebktB* resulted in an increase of lipid synthesis by 36%, with a constant ratio of OCFAs to total lipids. This increase of total lipids probably results from a similar increase of acetoacetyl-CoA (C4) and β-ketovaleryl-CoA (C5) due to the broad substrate specificity of β-ketothiolase [Mitsky et al. 1998], thus leading to constant ratio of OCFAs to total lipids.

## 5. Conclusion

In this work, we demonstrated the improvement of OCFA production by improving C3 and C5 precursor pools in *Y. lipolytica*. The overexpression of propionyl-CoA transferases (*Cppct* and *Repct*) or propionyl-CoA synthetase (*SeprpE*) in order to increase the propionyl-CoA precursor significantly improved the OCFAs synthesis, the best result was obtained with the overexpression of *Repct*. In *Repct*-expressing strain, the growth was negatively impacted in propionate medium, while it was restored and leads to a high OCFA ratio (53.2 % of total lipids) when complemented with acetate. We attributed the defect of growth to a limitation of the acetyl-CoA pool due to the transfer of its CoA moiety to propionyl-CoA. Our experiments also showed that increasing acetate had a negative effect on the OCFA ratio. As the synthesis of ECFA and OCFA are different only at the first condensation step of lipid synthesis depending on the primer, acetyl-CoA and propionyl-CoA, respectively, an excessive acetyl-CoA pool would beneficiate to ECFA production. This indicates a dual role of acetyl-CoA in OCFA production and the necessity of a fine-tuned balance between propionyl-CoA and acetyl-CoA for optimizing the production of OCFAs. Further engineering to accumulate higher amounts of lipids and provide C5 precursor, and optimization of carbon and nitrogen sources promoted OCFA production up to 1.87 g/L representing 62% of total lipids. This result shows the highest recombinant OCFA titer reported in yeast, to date. This study paves the way for future metabolic engineering strategies for the microbial production of OCFAs and its derivatives having high potentials in the pharmaceutical, food, and chemical industries.

## Abbreviations

OD: optical density
DCW: dry cell weight
FA: fatty acid
OCFA: odd-chain fatty acid
ECFA: even-chain fatty acid
FAME: fatty acid methyl ester
TAG: triacylglycerol
PDH: pyruvate dehydrogenase complex
PHBV: poly(3-hydroxybutyrate-co-3-hydroxyvalerate)

## Author contributions

**YKP:** Conceptualization, Data curation, Formal analysis, Investigation, Validation, Writing – original draft, Writing – review & editing. **FB:** Resources, Writing – review & editing, **FL:** Resources, Writing – review & editing. **JMN:** Conceptualization, Supervision, Writing – review & editing.

## Acknowledgments

We would like to acknowledge the Kwanjeong Educational Foundation (KEF) for the PhD scholarship of YKP.

