## Supplementary materials for "Engineering precursor pools for increasing production of odd-chain fatty acids in *Yarrowia lipolytica*"

**Supplementary Table 1.** The primers used in this study.

| Name | Sequence (5' – 3') |
|---|---|
| pTEF-internal-Fw | TCTGGAATCTACGCTTGTTCA |
| ACS2-Fw | ATGTCTGAAGACCACCCAGC |
| ACS2-intern-Rev | GCTCGTGGGTGTTCCACA |
| CpPCT-intern-Rev | GCAACAGAAGCCACGTACTCA |
| EnPCT-Fw | ATGACCCACCCCCAGCAG |
| EnPCT-internal-Rev | GACCGGCAGATCCAGGTCGG |
| EcPCT-Fw | ATGAAACCTGTCAAACCGCC |
| EcPCT-internal-Rev | GATTCCTTGCGAAGTGATTCCG |
| RePCT-Fw | ATGAAGGTGATTACCGCCAGAG |
| RePCT-internal-Rev | CCAATGGGGCCTGCCTC |
| EcPrpE-Fw | ATGTCTTTCTCCGAGTTCTACCAG |
| EcPrpE-Rev | CTACTCCTCCATAGCCTGTCG |
| SePrpE-Fw | ATGTCTTTCTCCGAGTTCTACCAG |
| SePrpE-Rev | CTACTCCTCAATGGCCTGTCG |
| ReBktB-Fw | ATGACCCGAGAGGTGGTGGTGGTC |
| ReBktB-internal-Rev | CCTTGAAGTAGCCGGCCTTGATGG |
| ReBktB-Rev | GATTCGCTCGAAGATGGCAGCGATG |

**Supplementary Table 2.** The sequences of the genes used in this study.

| Name | Sequence (5' – 3') |
|---|---|
| *Repct* | ATGAAGGTGATTACCGCCAGAGAAGCAGCGGCTCTTGTGCAGGACGGTTGGACTGTTGCATCGGCTGGATTCGT<br>TGGCGCAGGCCATGCTGAGGCAGTCACCGAAGCCCTTGAGCAGCGATTTCTGCAATCGGGTCTGCCACGAGATC<br>TGACCCTCGTCTACTCTGCTGGACAGGGCGATCGTGGTGCCCGAGGTGTGAACCACTTCGGCAATGCCGGCATG<br>ACCGCCAGCATCGTCGGCGGCCATTGGAGATCCGCAACCAGACTCGCCACCCTGGCCATGGCTGAGCAGTGTGA<br>GGGCTACAACCTGCCTCAAGGCGTCCTTACGCACCTGTACCGAGCCATTGCTGGCGGTAAACCTGGTGTCATGAC<br>CAAGATCGGCCTCCATACGTTCGTCGACCCACGAACCGCCCAAGATGCCCGATACCATGGCGGCGCGGTTAACG<br>AGCGAGCACGGCAGGCCATTGCCGAGGGAAAGGCTTGCTGGGTTGACGCCGTGGACTTTCGAGGCGATGAGTAC<br>CTGTTCTACCCCTCGTTTCCCATCCACTGTGCGCTCATTCGGTGCACTGCCGCTGACGCCCGAGGAAACCTCTCCA<br>CTCACAGAGAGGCCTTTCACCACGAACTTTTGGCAATGGCCCAAGCTGCTCACAACTCCGGAGGCATCGTCATCG<br>CGCAGGTGGAGTCCCTCGTGGACCACCACGAGATTCTGCAGGCCATCCACGTTCCAGGCATTCTGGTGGACTAC<br>GTCGTGGTTTGCGACAACCCCGCTAATCACCAGATGACCTTCGCCGAGTCCTACAACCCTGCGTACGTCACGCCT<br>TGGCAGGGAGAAGCTGCCGTGGCCGAAGCCGAGGCCGCTCCCGTCGCTGCTGGACCCCTTGACGCGCGGACCAT<br>CGTGCAGCGTCGAGCCGTTATGGAGCTGGCCCGACGAGCCCCGCGAGTTGTGAACCTCGGTGTCGGAATGCCTG<br>CTGCCGTTGGTATGCTCGCCCATCAGGCTGGACTCGACGGCTTCACCCTGACTGTGGAGGCAGGCCCCATTGGTG<br>GTACTCCCGCTGACGGACTGTCCTTTGGTGCCTCTGCTTATCCGGAGGCTGTCGTCGACCAGCCTGCCCAGTTCG<br>ACTTCTACGAAGGCGGTGGCATTGACCTTGCCATCCTCGGCTTGGCTGAGCTCGATGGTCACGGCAACGTCAACG<br>TGTCCAAGTTCGGTGAGGGAGAAGGAGCCTCCATTGCTGGTGTTGGCGGTTTCATCAACATCACCCAGTCTGCTC<br>GAGCCGTCGTGTTCATGGGAACACTGACAGCAGGTGGACTTGAGGTTCGAGCTGGTGATGGAGGACTCCAGATC<br>GTCCGAGAGGGCCGAGTCAAGAAGATCGTCCCTGAGGTGTCTCACCTGTCCTTTAACGGTCCCTATGTGGCTTCT<br>CTCGGAATCCCTGTCCTGTACATCACTGAGCGAGCTGTTTTCGAGATGCGAGCTGGAGCTGATGGCGAAGCCCG<br>ATTGACTCTGGTGGAGATTGCGCCCGGTGTCGACCTTCAGCGGGACGTTTTGGACCAGTGTAGCACACCCATTGC<br>TGTCGCCCAGGATCTGCGTGAGATGGATGCCCGTCTGTTTCAGGCCGGTCCCCTGCATCTGTAA |
| *Cppct* | ATGCGAAAGGTTCCCATCATCACTGCTGACGAGGCTGCCAAGCTCATCAAGGACGGAGATACCGTTACTACTTC<br>GGGTTTTGTCGGAAACGCTATCCCTGAGGCTCTGGACCGAGCTGTCGAGAAGCGATTCCTCGAGACCGGCGAGC<br>CTAAGAACATTACTTACGTTTACTGTGGATCTCAGGGTAACCGAGACGGACGAGGTGCTGAGCACTTTGCCCATG<br>AGGGCCTGCTCAAGCGATACATTGCTGGACACTGGGCCACCGTTCCCGCTCTGGGAAAGATGGCCATGGAGAAC<br>AAGATGGAGGCTTACAACGTGTCCCAGGGAGCCCTGTGCCACCTCTTCCGAGACATCGCCTCGCATAAGCCCGG<br>TGTTTTCACCAAGGTCGGCATCGGAACTTTTATTGACCCTCGAAACGGCGGCGGCAAGGTCAACGACATCACCA<br>AGGAAGACATTGTTGAGCTGGTGGAGATTAAGGGCCAGGAGTACCTCTTCTACCCCGCCTTTCCTATCCACGTGG<br>CTCTGATTCGAGGAACCTACGCCGACGAGTCCGGTAACATCACTTTTGAGAAGGAAGCCGCTCCCCTCGAGGGA<br>ACCTCTGTCTGTCAGGCTGTTAAGAACTCCGGCGGAATTGTGGTCGTTCAGGTCGAGCGAGTGGTCAAGGCCGG<br>AACTCTGGACCCCCGACATGTCAAGGTTCCTGGTATCTACGTGGATTACGTTGTGGTCGCTGACCCCGAGGATCA<br>CCAGCAGTCGCTGGACTGCGAGTACGATCCCGCCCTCTCTGGCGAGCATCGACGACCTGAGGTTGTGGGAGAGC<br>CCCTGCCTCTCTCGGCTAAGAAGGTCATCGGCCGACGAGGAGCCATTGAGCTGGAGAAGGACGTGGCTGTCAAC<br>CTCGGTGTGGGAGCTCCTGAGTACGTGGCTTCTGTTGCTGACGAGGAAGGCATCGTCGATTTCATGACCCTGACT<br>GCCGAGTCTGGAGCTATTGGTGGCGTGCCTGCTGGAGGTGTCCGATTCGGTGCCTCCTACAACGCCGACGCTCTG<br>ATTGATCAGGGCTACCAGTTTGACTACTACGATGGCGGAGGTCTGGACCTCTGTTACCTGGGTCTCGCTGAGTGC<br>GATGAGAAGGGCAACATCAACGTGTCCCGATTCGGTCCCCGAATTGCCGGCTGTGGCGGCTTCATCAACATTAC<br>CCAGAACACTCCTAAGGTTTTCTTTTGCGGCACCTTCACTGCTGGTGGCCTGAAGGTGAAGATCGAGGACGGCAA<br>GGTCATCATTGTCCAGGAAGGCAAGCAGAAGAAGTTCCTGAAGGCCGTCGAGCAGATCACCTTTAACGGAGACG<br>TTGCCCTCGCTAACAAGCAGCAGGTGACCTACATTACTGAGCGATGTGTCTTCCTGCTCAAGGAAGACGGTCTGC<br>ACCTCTCTGAGATTGCTCCTGGCATTGATCTGCAGACCCAGATCCTCGACGTGATGGATTTTGCTCCTATCATTGA<br>CCGAGATGCCAACGGCCAGATTAAGCTGATGGATGCTGCTCTGTTTGCTGAGGGTCTGATGGGCCTGAAGGAGA<br>TGAAGTCTTAG |
| *Ecpct* | ATGAAACCTGTCAAACCGCCTCGAATCAACGGCCGAGTTCCAGTTCTCTCTGCCCAGGAAGCCGTTAACTACATT<br>CCCGATGAGGCTACCCTCTGTGTCCTTGGCGCTGGAGGAGGCATTCTTGAGGCCACCACGCTGATTACAGCCCTG<br>GCTGACAAGTACAAGCAGACGCAGACTCCCCGAAATCTGTCCATTATCTCTCCCACAGGACTTGGTGATCGAGCT<br>GATCGAGGCATTTCCCCTCTGGCACAAGAGGGACTGGTGAAGTGGGCGCTGTGCGGTCATTGGGGCCAGTCTCC<br>ACGAATTAGCGATCTGGCCGAACAGAACAAGATTATTGCCTACAACTACCCTCAGGGTGTGCTTACCCAGACCC<br>TCCGAGCCGCAGCTGCCCATCAACCCGGCATTATCTCCGACATCGGCATTGGAACCTTTGTCGATCCCCGACAGC<br>AGGGCGGCAAGCTGAACGAGGTGACCAAAGAGGACCTCATCAAGTTGGTTGAGTTCGACAACAAGGAGTACCT<br>TTACTACAAGGCCATTGCTCCCGATATTGCCTTCATTCGTGCAACCACCTGCGATTCCGAAGGCTACGCCACTTTT<br>GAGGACGAGGTGATGTATCTCGACGCCCTGGTTATTGCGCAAGCTGTCCACAACAACGGTGGAATCGTGATGAT<br>GCAGGTCCAGAAGATGGTTAAGAAGGCCACGCTTCACCCCAAGTCCGTGCGTATCCCCGGTTACCTCGTCGACA<br>TCGTGGTCGTTGACCCGGATCAGTCTCAGTTGTATGGTGGCGCCCCAGTCAACCGATTCATCTCTGGCGACTTCA<br>CCCTCGACGACTCCACCAAGCTGTCGCTTCCCCTCAATCAGCGGAAGCTTGTCGCTAGACGAGCACTGTTTGAGA<br>TGCGGAAAGGAGCGGTCGGAAACGTGGGTGTCGGCATTGCCGATGGTATCGGACTCGTTGCCCGAGAAGAAGGT<br>TGTGCTGACGACTTCATTTTGACCGTCGAGACTGGCCCTATCGGCGGAATCACTTCGCAAGGAATCGCCTTTGGC<br>GCCAATGTCAACACCCGAGCCATCCTTGACATGACGTCCCAGTTTGACTTCTACCACGGAGGAGGTCTGGACGTG<br>TGCTACCTGTCGTTTGCAGAAGTCGACCAGCATGGCAACGTTGGTGTCCACAAGTTCAACGGCAAGATCATGGG<br>AACCGGAGGCTTCATCGACATCTCCGCTACTTCCAAGAAGATCATCTTCTGTGGCACACTCACCGCTGGTTCTCT<br>CAAGACTGAGATTGCTGACGGTAAGCTGAACATTGTGCAGGAGGGCCGAGTCAAGAAGTTCATCCGAGAACTGC<br>CTGAGATCACCTTCAGCGGCAAGATCGCCCTGGAGAGAGGTCTGGATGTGCGGTACATCACAGAGAGAGCTGTG<br>TTTACTCTGAAAGAGGATGGTCTGCACTTGATCGAGATTGCTCCTGGTGTTGACCTGCAGAAGGACATCCTCGAC<br>AAGATGGATTTCACTCCCGTGATCTCCCCTGAGCTGAAGCTGATGGACGAGCGACTCTTCATTGACGCTGCCATG<br>GGTTTTGTCCTCCCCGAGGCTGCGCACTAA |

| | |
|---|---|
| *Enpct* | ATGACCCACCCCCAGCAGGCCGTTCACGCCGCTTCGCTCCAGAACCCCGAGGCTTTTTGGTCCCATCACGCCCAG<br>CAGCTCCATTGGCACAAGAAGCCCTCGCGAGCCATTGGCCGATCTACCAAGACTCTGGCTTCTGGAGCCTCCCAC<br>GAGTCCTGGTCGTGGTTCCCTGACGGAGAGATCTCCACCACTTACAACTGTGTGGATCGACATGTCCTGAACGGC<br>AACGGAGACAACGTGGCCATCATTTGGGATTCTGCTGTCACCGGCAAGAAGGAGAAGTACACTTACCGACAGCT<br>GCTCGACGAGGTCGAGGTTCTGGCTGGTGTCCTCCGAGAGGAGGGCGTTAAGAAGGGAGACGTGGTCATCATCT<br>ACATGCCCATGATCCCTGCCGCTCTGATTGGAGCTCTCGCTGTCGCTCGACTGGGTGCTATTCACGCCGCTGTTTT<br>CGGCGGATTTGCCGCTAAGTCCCTGGCTCAGCGAATTGAGGCTGCTCGACCCCGAGCTATCCTCACCGCTTCTTG<br>CGGTATTGAGGGCGCCAAGGGACCCATCGCTTACCGACCTCTGGTGGAGGGCGCTATTGAGGCCTCTTCCTTCAA<br>GCCCGAGAAGGTCCTGATCTGGCAGCGAGACCAGCTCCGATGGAACAACCCTGATAAGCTGGGTGGCCAGCGA<br>AACTGGAACCGACTCGTGAAGTCCGCCCGAATGCGAGGCATTCGAGCTGAGCCCGTGCCTGTCCGATCTACCGA<br>CGGACTGTACATCATCTACACTTCCGGTACCACTGGCCTCCCCAAGGGAGTTGTGCGAGAGGCCGGAGGTCACG<br>CTGTGGGTCTGTCTCTCTCCATCAAGTACCTGTTCGACATTCATGGTCCCGGCGATACCATGTTTTGTGCCTCCGA<br>CATTGGTTGGGTCGTTGGCCACTCGTACATCCTGTACGCCCCTCTGCTCGTCGGAGCTACCACTGTTCTCTTCGAG<br>GGAAAGCCTGTGGGTACCCCTGACGCTGGTACTTTTTGGCGAGTGGTCGCCGAGCATAAGGCTAACGTCCTGTTC<br>ACCGCTCCCACTGCCCTCCGAGCTATTCGAAAGGAGGACCCTGATAACAAGCACTTTGAGAAGGTGGCCGGTGA<br>CAACAACCTGCGACATCTCCGAGCCCTGTTCCTCGCTGGCGAGCGATCGGAGCCCTCTATCGTCCGAGCCTACCA<br>GGACCTGCTCACCAAGCACGCCGCTCGAGGAGCTCTGGTTGTGGATAACTGGTGGTCGTCTGAGTCGGGCTCTCC<br>TATTTCCGGACTGGCTCTCCGATCGGCTGTCGGTCGAGTTCCTCCTCGATCGGACGAGTACGATGTGGCCCCCCT<br>GGCTATCCGACCTGGATCTGCCGGTCTCCCCATGCCTGGTTTCGACGTCCGAGTCGTTGACGATGAGGGCAACGA<br>GGTTGCCCAGGGCACCATGGGAAACATTGTGATGGCTACTCCCCTGGCCCCTACCGCTTTCACTCGACTCTTTAA<br>CGACGATGAGCGATTCTACAAGGGATACCTGAAGCGATTTGGCGGACGATGGCTCGACACCGGCGACGCTGGTA<br>TGATCGACCAGGATGGCTACATTCACGTGATGTCCCGATCGGACGATATCATTAACGTCGCCGCTCACCGATTCT<br>CTACTGGACAGGGTTCCATCGAGCAGGCCATTCTGTCGCACCCCGCCATTGGAGAGGCTTCTGTGGTCGGCATCC<br>CCGACGCCCTGAAGGGACATCTCCCTTTCGCTTTTATCACCCTGAAGCAGTCCGGTGGTAACTCGCCTGCTCGAC<br>CTTCTGCTGAGCTGTTCAACTCCGTTAACCGACTCGTGCGAGAGCAGATCGGAGCTATTGCCTCCCTGGGAGGAA<br>TGATCCAGGGCCAGGGAATGATTCCCAAGACCCGATCTGGCAAGACTCTCCGACGAGTGCTGCGAGAGCTCGTC<br>GAGAACGGAGCCCGAGGTGAGTTCGAGAAGGAGGTTGCTGTGCCTCCTACCGTGGAGGACCGAGGCGTTGTGG<br>AGGTTGCCCGAGAGAAGGTGCGAGAGTACTTCGAGTCTCAGTCCGGATCGCCCAAGGCTAAGCTGTAG |
| *EcprpE* | ATGTCTTTCTCCGAGTTCTACCAGCGATCTATCAACGAGCCTGAGCAGTTCTGGGCTGAGCAGGCTCGACGAATT<br>GACTGGCAGACCCCCTTCACCCAGACCCTGGACCACTCCAACCCTCCCTTCGCCCGATGGTTCTGTGAGGGCCGA<br>ACCAACCTGTGCCACAACGCTATCGACCGATGGCTGGAGAAGCAGCCTGAGGCTCTGGCTCTGATTGCCGTCTCT<br>TCCGAGACTGAGGAAGAGCGAACCTTCACCTTCCGACAGCTGCACGACGAGGTGAACGCCGTCGCTTCTATGCT<br>GCGATCCCTGGGAGTGCAGCGAGGTGACCGAGTGCTGGTCTACATGCCCATGATCGCCGAGGCTCACATTACCC<br>TGCTGGCCTGTGCTCGAATCGGTGCCATTCACTCTGTGGTCTTCGGCGGATTCGCTTCTCACTCCGTGGCCGCTCG<br>AATCGACGACGCCAAGCCCGTGCTGATTGTGTCCGCTGACGCTGGAGCTCGAGGTGGCAAGATCATTCCCTACA<br>AGAAGCTGCTGGACGACGCTATCTCTCAGGCTCAGCACCAGCCCCGACACGTGCTGCTGGTGGACCGAGGCCTG<br>GCTAAGATGGCTCGAGTGTCTGGACGAGACGTCGACTTCGCTTCTCTGCGACACCAGCACATTGGTGCTCGAGTG<br>CCTGTGGCTTGGCTGGAGTCTAACGAGACTTCCTGCATTCTGTACACCTCTGGTACCACCGGCAAGCCCAAGGGA<br>GTGCAGCGAGACGTCGGAGGTTACGCTGTCGCCCTGGCTACCTCCATGGACACCATTTTCGGCGGAAAGGCCGG<br>ATCTGTGTTCTTCTGTGCTTCCGACATCGGCTGGGTGGTCGGACACTCCTACATTGTCTACGCTCCCCTGCTGGCC<br>GGCATGGCTACCATCGTGTACGAGGGACTGCCCACCTGGCCTGACTGTGGTGTGTGGTGGACCATTGTCGAGAA<br>GTACCAGGTGTCTCGAATGTTCTCCGCCCCCACCGCTATCCGAGTGCTGAAGAAGTTCCCCACCGCCGAGATTCG<br>AAAGCACGACCTGTCTTCCCTGGAGGTCCTGTACCTGGCTGGAGAGCCTCTGGACGAGCCTACCGCTTCTTGGGT<br>GTCCAACACCCTGGACGTGCCCGTCATCGACAACTACTGGCAGACCGAGTCTGGTTGGCCCATCATGGCCATTGC<br>TCGAGGCCTGGACGACCGACCTACCCGACTGGGCTCTCCCGGTGTCCCCATGTACGGATACAACGTCCAGCTGCT<br>GAACGAGGTGACCGGAGAGCCCTGTGGCGTGAACGAGAAGGGTATGCTGGTGGTGGAGGGTCCTCTGCCTCCCG<br>GTTGCATCCAGACCATTTGGGGCGACGACGGACGATTCGTCAAGACCTACTGGTCCCTGTTCTCTCGACCCGTGT<br>ACGCCACCTTCGACTGGGGAATCCGAGACGCTGACGGTTACCACTTCATCCTGGGCCGAACCGACGACGTGATT<br>AACGTCGCCGGACACCGACTGGGTACCCGAGAGATCGAGGAGTCTATTTCTTCTCACCCTGGAGTGGCTGAGGT<br>GGCTGTGGTCGGAGTCAAGGACGCTCTGAAGGGTCAGGTGGCCGTCGCTTTCGTGATTCCCAAGGAGTCTGACT<br>CCCTGGAGGACCGAGACGTCGCCCACTCTCAGGAGAAGGCCATCATGGCTCTGGTGGACTCCCAGATTGGTAAC<br>TTCGGCCGACCCGCTCACGTGTGGTTCGTCTCTCAGCTGCCCAAGACCCGATCCGGAAAGATGCTGCGACGAACC<br>ATCCAGGCCATTTGTGAGGGTCGAGATCCCGGCGACCTGACCACCATCGACGACCCTGCTTCCCTGGACCAGATT<br>CGACAGGCTATGGAGGAGTAG |
| *SeprpE* | ATGTCTTTCTCCGAGTTCTACCAGCGATCTATCAACGAGCCTGAGGCTTTCTGGGCTGAGCAGGCTCGACGAATT<br>GACTGGCGACAGCCCTTCACCCAGACCCTGGACCACTCTCGACCTCCCTTCGCTCGATGGTTCTGTGGCGGAACC<br>ACCAACCTGTGCCACAACGCCGTCGACCGATGGCGAGACAAGCAGCCTGAGGCTCTGGCTCTGATTGCCGTGTC<br>TTCCGAGACTGACGAGGAGCGAACCTTCACCTTCTCTCAGCTGCACGACGAGGTGAACATTGTCGCCGCTATGCT<br>GCTGTCCCTGGGCGTGCAGCGAGGCGACCGAGTGCTGGTCTACATGCCCATGATCGCCGAGGCTCAGATTACCC<br>TGCTGGCCTGTGCTCGAATCGGCGCTATTCACTCTGTGGTCTTCGGTGGCTTCGCTTCTCACTCTGTGGCTGCTCG<br>AATCGACGACGCTCGACCCGCCCTGATTGTGTCTGCTGACGCTGGAGCTCGAGGCGGCAAGATCCTGCCCTACA<br>AGAAGCTGCTGGACGACGCTATTGCCCAGGCTCAGCACCAGCCCAAGCACGTGCTGCTGGTCGACCGAGGTCTG<br>GCTAAGATGGCCTGGGTGGACGGCCGAGATCTGGACTTCGCTACCCTGCGACAGCAGCACCTGGGTGCTTCTGT<br>GCCCGTCGCCTGGCTGGAGTCTAACGAGACTTCCTGCATCCTGTACACCTCCGGCACCACCGGAAAGCCTAAGG<br>GAGTGCAGCGAGACGTCGGCGGATACGCTGTGGCTCTGGCTACCTCTATGGACACCATTTTCGGTGGCAAGGCT<br>GGAGGTGTGTTCTTCTGTGCCTCTGACATCGGTTGGGTGGTCGGCCACTCCTACATTGTCTACGCTCCCCTGCTGG<br>CTGGTATGGCCACCATCGTGTACGAGGGCCTGCCTACCTACCCTGACTGCGGAGTGTGGTGGAAGATTGTCGAG<br>AAGTACCAGGTGAACCGAATGTTCTCCGCTCCCACTGCCATCCGAGTCCTGAAGAAGTTCCCCACCGCCCAGATT<br>CGAAACCACGACCTGTCTTCTCTGGAGGCTCTGTACCTGGCTGGAGAGCCTCTGGACGAGCCTACCGCCTCTTGG<br>GTGACCGAGACTCTGGGCGTGCCCGTCATCGACAACTACTGGCAGACCGAGTCCGGTTGGCCCATTATGGCTCTG<br>GCTCGAGCTCTGGACGACCGACCTTCCCGACTGGGTTCCCCCGGAGTCCCTATGTACGGATACAACGTCCAGCTG<br>CTGAACGAGGTGACCGGAGAGCCCTGTGGTATCAACGAGAAGGGCATGCTGGTCATCGAGGGACCTCTGCCTCC<br>CGGTTGCATCCAGACCATTTGGGGAGACGACGCCCGATTCGTCAAGACCTACTGGTCCCTGTTCAACCGACAGGT<br>GTACGCTACCTTCGACTGGGGCATCCGAGACGCCGAGGGATACTACTTCATTCTGGGTCGAACCGACGACGTGA<br>TCAACATTGCCGGCCACCGACTGGGAACCCGAGAGATCGAGGAGTCTATTTCTTCCTACCCTAACGTGGCTGAG |

| | |
|---|---|
| | GTGGCTGTGGTCGGCATCAAGGACGCTCTGAAGGGACAGGTGGCCGTCGCTTTCGTCATTCCCAAGCAGTCTGA<br>CACCCTGGCTGACCGAGAGGCTGCTCGAGACGAGGAGAACGCTATCATGGCCCTGGTGGACAACCAGATTGGAC<br>ACTTCGGTCGACCCGCTCACGTGTGGTTCGTCTCTCAGCTGCCCAAGACCCGATCCGGCAAGATGCTGCGACGAA<br>CCATCCAGGCCATTTGTGAGGGTCGAGATCCCGGCGACCTGACCACCATCGACGACCCTGCTTCCCTGCAGCAG<br>ATCCGACAGGCCATTGAGGAGTAG |
| *YlACS2* | ATGTCTGAAGACCACCCAGCCATCCACCCACCCTCCGAGTTCAAGGACAACCACCCCCACTTCGGAGGCCCCCA<br>CCTCGACTGTCTGCAGGACTACCACCAGCTGCACAAGGAGTCCATTGAGGACCCCAAGGCCTTCTGGAAGAAGA<br>TGGCCAACGAGCTCATCTCCTGGTCAACCCCCTTTGAAACTGTGCGATCTGGCGGCTTCGAGCACGGCGACGTGG<br>CCTGGTTCCCCGAGGGCCAGCTCAACGCCTCCTACAACTGTGTGGATCGACACGCCTTTGCCAACCCCGACAAGC<br>CCGCCATCATTTTTGAGGCCGATGAGCCGGGCCAGGGCCGAATCGTCACCTACGGCGAACTGCTGCGACAGGTG<br>TCTCAGGTCGCAGCCACCCTGCGATCCTTCGGCGTCCAGAAGGGCGATACTGTGGCCGTCTACCTGCCCATGATC<br>CCCGAGGCCATTGTCACTCTGCTGGCCATCACCCGAATTGGCGCTGTCCACTCGGTCATCTTCGCCGGCTTCTCCT<br>CCGGTTCTCTGCGAGACCGAATCAACGACGCCAAGTCCAAGGTTGTCGTCACCACCGACGCCTCCATGCGAGGA<br>GGCAAGACCATCGACACCAAGAAGATTGTCGATGAAGCCTTGCGAGACTGCCCCTCTGTTACCCACACCCTGGT<br>CTTCCGACGAGCAGGTGTCGAGAACCTGGCCTGGACTGAGGGCCGGGACTTCTGGTGGCACGAGGAGGTCGTCA<br>AGCACCGACCCTACCTTGCCCCCGTCCCCGTTGCCTCCGAGGACCCCATCTTCCTGCTTTACACCTCTGGATCCAC<br>CGGCACCCCCAAGGGTCTGGCCCACGCTACCGGTGGCTACCTGCTTGGTGCTGCCCTGACCGCCAAGTACGTGTT<br>TGACATCCACGGAGACGACAAGCTGTTCACCGCTGGAGACGTTGGCTGGATCACCGGCCACACCTACGTGCTCT<br>ACGGTCCTCTGATGCTCGGAGCCACCACTGTTGTGTTCGAGGGAACCCCTGCCTACCCCTCCTTCTCGCGATACT<br>GGGACATTGTCGACGACCACAAGATCACCCACTTCTACGTGGCTCCCACCGCCCTGCGTCTCCTGAAGCGGGCCG<br>GCACCCATCACATTAAGCACGACCTGTCGTCTCTGCGAACCCTCGGCTCTGTGGGTGAGCCCATTGCCCCCGACG<br>TGTGGCAGTGGTACAACGACAACATTGGCCGAGGCAAGGCCCACATCTGTGACACCTACTGGCAGACCGAGACT<br>GGCTCGCATATCATTGCCCCCATGGCCGGCGTGACCCCCACCAAGCCCGGTTCTGCTTCCCTGCCTGTCTTTGGA<br>ATTGATCCCGTTATCATTGATCCCGTGTCTGGCGAGGAGCTCAAGGGTAACAACGTTGAGGGTGTTCTTGCCCTG<br>CGATCTCCCTGGCCCTCCATGGCCCGAACCGTGTGGAACACCCACGAGCGATACATGGAGACCTACCTGCGGCC<br>CTACCCCGGCTACTACTTCACCGGTGATGGTGCTGCCCGAGACAATGACGGCTTTTACTGGATCCGAGGCCGAGT<br>CGACGACGTTGTCAACGTTTCTGGCCACCGTCTTTCCACCGCCGAGATTGAGGCTGCTCTCATTGAGCACGCTCA<br>GGTGTCTGAGTCTGCCGTTGTTGGTGTCCATGACGATCTGACTGGCCAGGCCGTCAACGCCTTTGTGGCTCTCAA<br>GAACCCCGTCGAGGATGTGGACGCTCTGCGAAAGGAGCTTGTTGTGCAGGTGCGAAAGACCATTGGACCCTTTG<br>CTGCTCCCAAGAATGTCATCATCGTGGACGATCTGCCCAAGACTCGGTCTGGCAAGATCATGCGACGAATTCTGC<br>GAAAGGTGCTTGCTGGCGAGGAGGACCAGCTCGGAGACATTTCCACTCTTGCTAACCCCGACGTTGTCCAGACC<br>ATCATTGAGGTTGTTCACTCGTTGAAAAAGTAA |
| *RebktB* | ATGACCCGAGAGGTGGTGGTGGTCAGCGGTGTTCGAACCGCCATCGGCACCTTCGGCGGCTCCCTGAAGGACGT<br>CGCCCCCGCTGAGCTGGGCGCTCTGGTTGTCCGAGAGGCCCTGGCCCGAGCTCAGGTGTCTGGTGACGACGTGG<br>GCCACGTCGTCTTCGGCAACGTCATTCAGACCGAGCCCCGAGACATGTACCTGGGACGAGTCGCTGCCGTCAAC<br>GGCGGCGTCACCATTAACGCCCCCGCCCTCACCGTGAACCGACTGTGCGGTTCCGGCCTGCAGGCCATCGTCTCC<br>GCCGCTCAGACCATCCTGCTCGGCGACACCGACGTTGCCATCGGCGGCGGTGCCGAGTCCATGTCCCGAGCTCCC<br>TACCTGGCCCCCGCCGCTCGATGGGGTGCTCGAATGGGTGACGCCGGTCTGGTTGATATGATGCTGGGAGCCCTG<br>CACGACCCTTTCCACCGAATTCACATGGGCGTTACCGCTGAGAACGTCGCCAAGGAGTACGACATCTCCCGAGC<br>CCAGCAGGACGAGGCTGCCCTGGAGTCCCACCGACGAGCCTCCGCTGCCATCAAGGCCGGCTACTTCAAGGACC<br>AGATCGTCCCCGTCGTTTCCAAGGGCCGAAAGGGCGACGTCACCTTCGACACCGACGAGCACGTCCGACACGAC<br>GCCACCATCGACGACATGACCAAGCTGCGACCCGTCTTCGTCAAGGAGAACGGCACCGTGACCGCCGGAAACGC<br>CTCCGGACTGAACGACGCCGCCGCCGCTGTCGTGATGATGGAGCGAGCCGAGGCCGAGCGACGAGGACTGAAG<br>CCCCTGGCCCGACTGGTCAGCTATGGTCACGCCGGTGTGGACCCCAAGGCTATGGGAATTGGTCCCGTCCCCGCC<br>ACCAAGATCGCCCTGGAGCGAGCCGGACTGCAGGTGTCCGACCTGGACGTTATCGAGGCCAACGAGGCTTTCGC<br>TGCTCAGGCCTGCGCCGTTACCAAGGCCCTGGGCCTGGACCCCGCCAAGGTCAACCCCAACGGATCTGGCATCT<br>CCCTGGGCCACCCCATCGGCGCTACCGGTGCTCTGATCACCGTCAAGGCCCTGCACGAGCTCAACCGAGTCCAG<br>GGTCGATACGCCCTGGTCACCATGTGCATCGGAGGCGGCCAGGGCATCGCTGCCATCTTCGAGCGAATCTAA |

**Supplementary Table 3.** Production of OCFAs in obese-L strain with or without acetate. The strains were cultivated in YNBD2P0.5 and YNBD2P0.5A1 medium for 120 hours. Averages and standard deviations were obtained from two replicate experiments.

| Media | Lipid content % (g/g DCW) | | OCFA/total lipids (%) |
|---|---|---|---|
| | Total lipids | OCFAs | |
| YNBD2P0.5 | 18.81 ± 1.03 | 5.48 ± 1.05 | 28.94 ± 4.01 |
| YNBD2P0.5A1 | 17.27 ± 0.62 | 0.65 ± 0.01 | 3.79 ± 0.09 |

**Supplementary Table 4.** Production of OCFAs in yeasts.

| | Substrate (g/L) | Biomass (g/L) | Lipid (g/L) | Lipid content % (w/w) | OCFAs /Total lipids (%) | OCFAs (g/L) | Reference |
|---|---|---|---|---|---|---|---|
| *Trichosporon cutaneum* | Propionate 4 g/L | 1.60 | 0.38 | 23.90 | ~35 | 0.13 | Kolouchova et al. 2015 |
| *Trichosporon cutaneum* | Glucose 20 g/L<br>Propionate 4 g/L | 1.76 | 0.64 | 36.10 | < 20 | < 0.13 | Kolouchova et al. 2015 |
| *Cryptococcus curvatus* | Propionate 16.5 g/L | 6.7 | 2.28 | 34.1 | 38.7 | 0.88 | Zheng et al. 2012 |
| *Cryptococcus curvatus* | VFA 30 g/L<br>(A:P:B = 5:15:10*) | 8.24 | 3.83 | 46.47 | 42.3 | 1.37 | Liu et al. 2017 |
| *Yarrowia lipolytica* | Propionate 4 g/L | 3.53 | 0.31 | 8.90 | ~30 | <0.093 | Koluchova et al. 2015 |
| *Yarrowia lipolytica* | Glucose 20 g/L<br>Propionate 4 g/L | 3.83 | 0.39 | 10.20 | <15 | <0.040 | Koluchova et al. 2015 |
| *Yarrowia lipolytica* | Pentadecane 3 g/L<br>Rhamnolipids | 2.75 | 0.47 | 17.20 | 44.5 | 0.209 | Matatkova et al. 2017 |
| *Yarrowia lipolytica* (Obese) | Glucose 14 g/L<br>Propionate 4 g/L | 5.53 | 1.36 | 24.76 | 41.9 | 0.57 | Park et al. 2018 |
| *Yarrowia lipolytica* (Obese-LPB) | Glucose 20 g/L<br>Propionate 5 g/L<br>Acetate 10 g/L | 7.32 | 2.04 | 27.84 | 64.54 | 1.32 | This study |
| *Yarrowia lipolytica* (Obese-LPB) | Glucose 40 g/L<br>Propionate 10 g/L<br>Acetate 20 g/L | 10.52 | 3.01 | 28.56 | 62.07 | 1.87 | This study |

*A, acetate; P, propionate; B, Butyrate.

**Supplementary Figure 1.** Comparison of growth in wild-type and strains overexpressing propionate activating genes in (a) YNBD0.5, (b) YNBP0.5, (c) YNBD0.5P1, and (d) YNBP1A0.5. Averages were obtained from two replicate experiments.

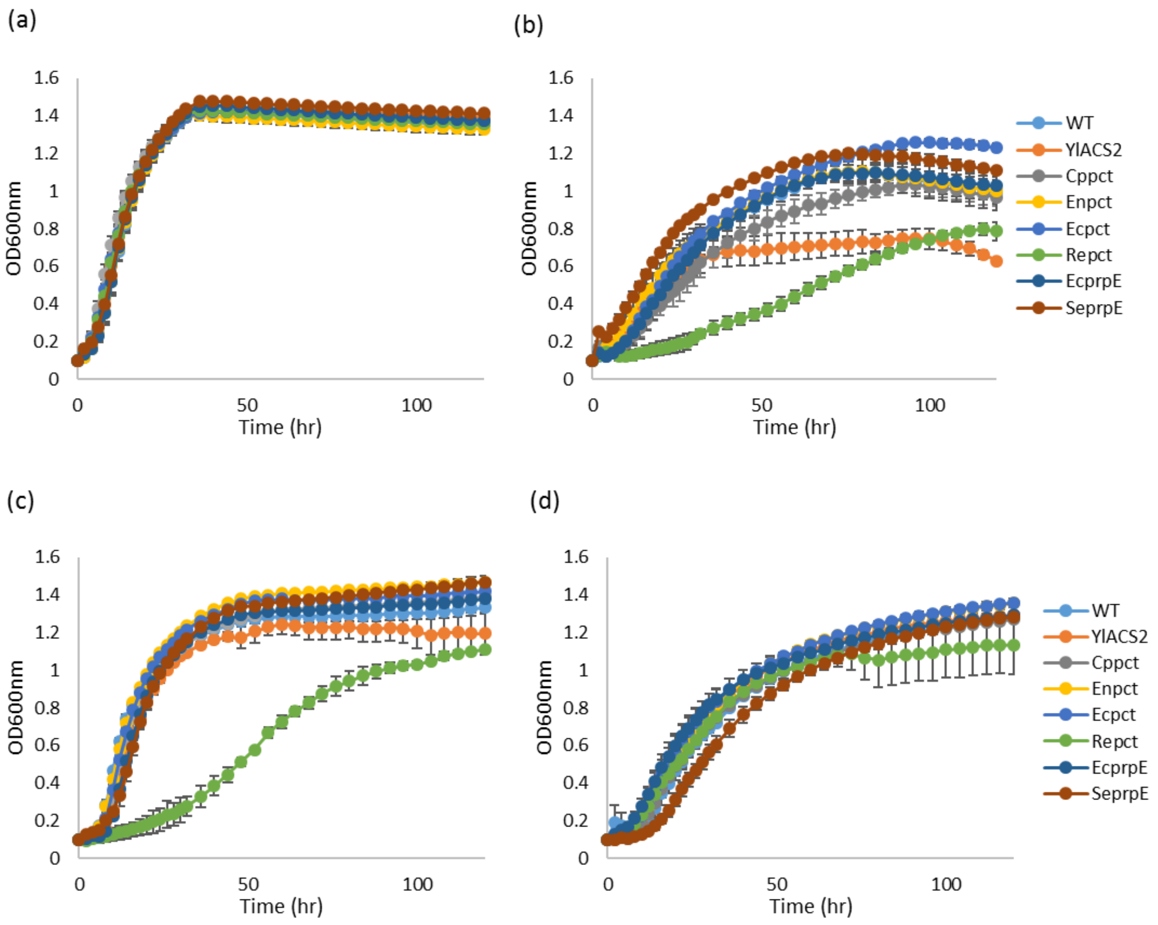

**Supplementary Figure 2.** Comparison of growth on different substrates in each strain overexpressing propionate activating genes. ●, YNBD0.5; ●, YNBP0.5; ●, YNBD0.5P1; ●, YNBP1A0.5. Averages were obtained from two replicate experiments.

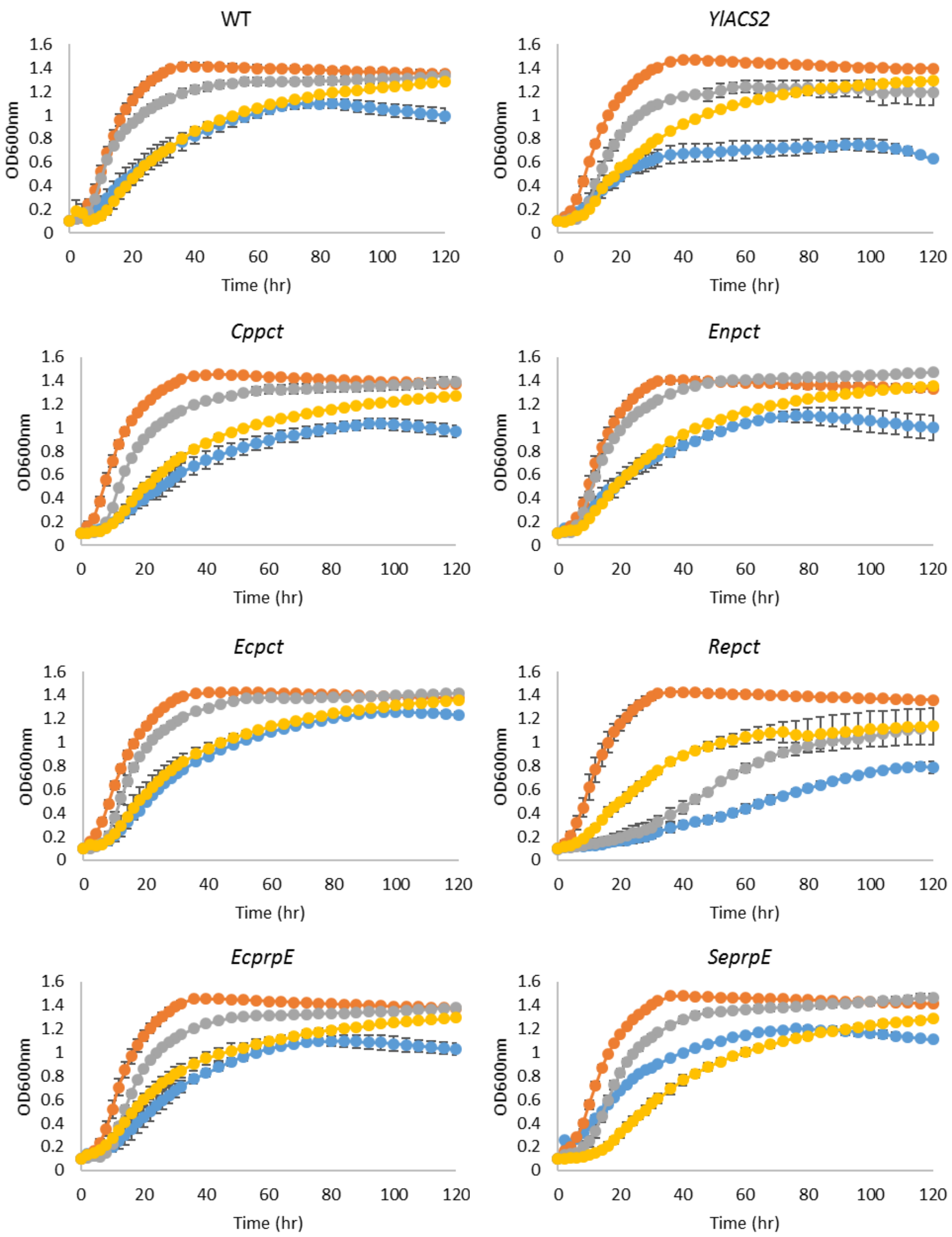

**Supplementary Figure 3.** (a) Lipid profiles (% in total lipids) of obese-L and obese-LP strains, (b) GC chromatogram of obese-L strain (JMY7228, control) and obese-LP (*Repct*) strain (JMY7780). The strains were cultivated in YNBD2P0.5A1 medium for 120 hours. Averages and standard deviations were obtained from two replicate experiments.

(a)

(b)

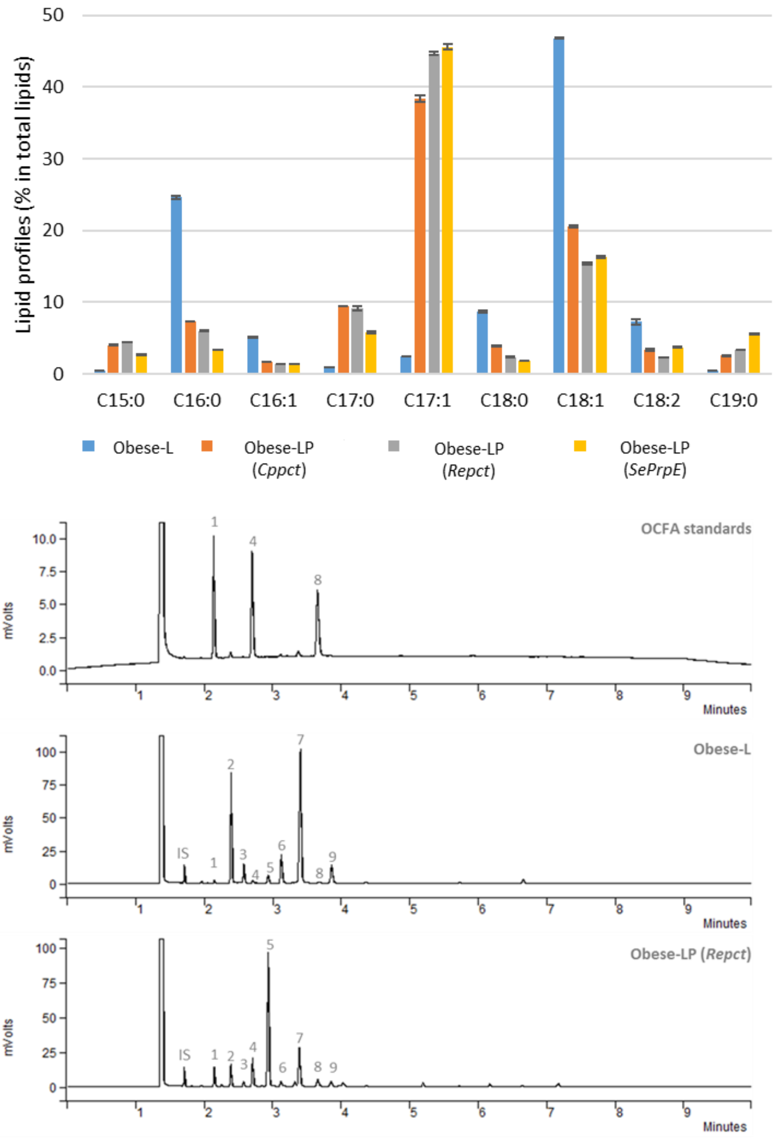


Peak list

IS - internal standard (C12:0, 100 µg)

1 – C15:0, 2 – C16:0, 3 – C16:1, 4 – C17:0, 5 – C17:1, 6 – C18:0, 7 – C18:1, 8 – C19:0, 9 – C18:2

**Supplementary Figure 4.** Substrate consumption of obese strains; (a) control, obese-L (b) obese-LP (*Cppct*), (c) obese-LP (*Repct*), and (d) obese-LP (*SeprpE*). Averages and standard deviations were obtained from two replicate experiments.

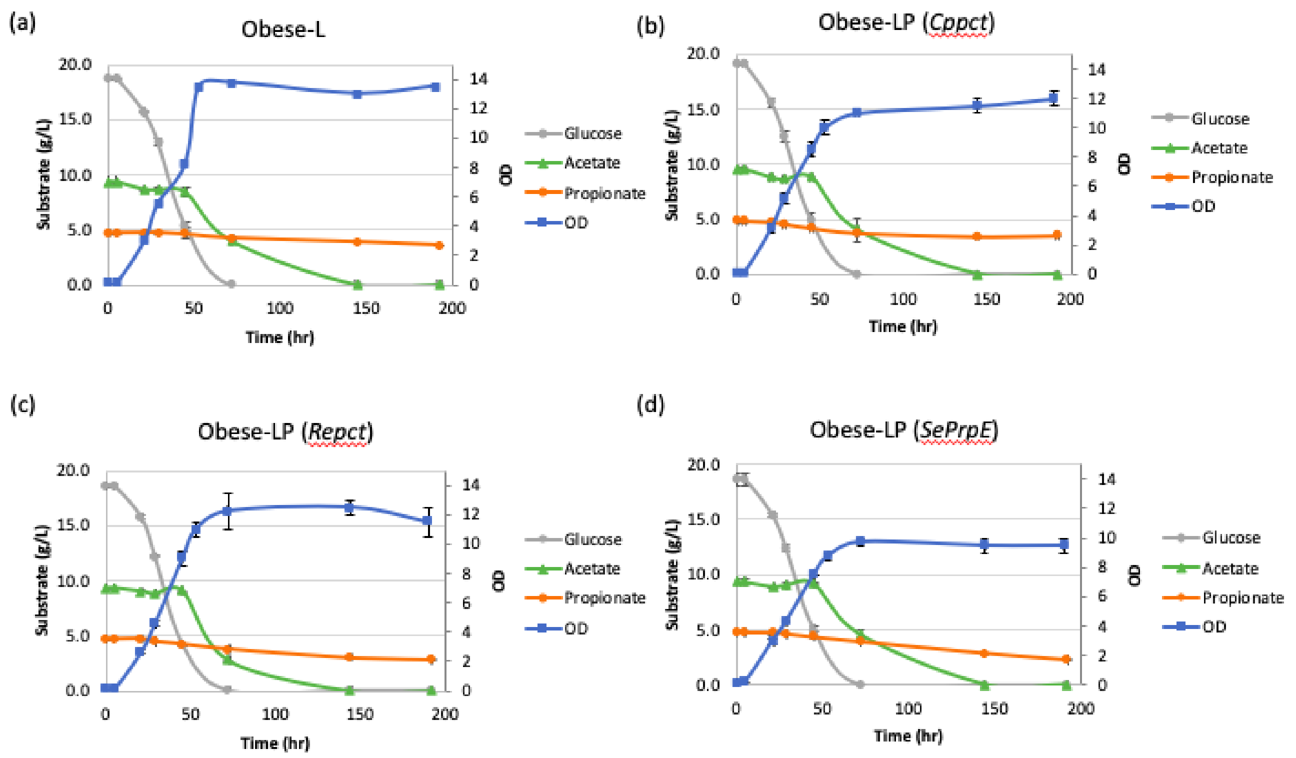
